# Human Myosin 1e tail but not motor domain replaces fission yeast Myo1 domains to support myosin-I function during endocytosis

**DOI:** 10.1101/535765

**Authors:** Sarah R. Barger, Michael L. James, Christopher D. Pellenz, Mira Krendel, Vladimir Sirotkin

**Affiliations:** From the Department of Cell and Developmental Biology, State University of New York Upstate Medical University, Syracuse, NY 13210

**Keywords:** endocytosis, myosin, actin, yeast, calmodulin, myosin-I, myosin 1e, Myo1

## Abstract

In both unicellular and multicellular organisms, long-tailed class I myosins function in clathrin-mediated endocytosis. Myosin 1e (Myo1e) in vertebrates and Myo1 in fission yeast have similar domain organization, yet whether these proteins or their individual protein domains are functionally interchangeable remains unknown. In an effort to assess functional conservation of class I myosins, we tested whether human Myo1e could replace Myo1 in fission yeast *Schizosaccharomyces pombe* and found that it was unable to substitute for yeast Myo1. To determine if any individual protein domain is responsible for the inability of Myo1e to function in yeast, we created human-yeast myosin-I chimeras. By functionally testing these chimeric myosins *in vivo,* we concluded that the Myo1e motor domain is unable to function in yeast, even when combined with the yeast Myo1 tail and a full complement of yeast regulatory light chains. Conversely, the Myo1e tail, when attached to the yeast Myo1 motor domain, supports localization to actin patches and partially rescues the endocytosis defect in *myo1Δ* cells. Further dissection showed that both the TH1 and TH2-SH3 domains in the human Myo1e tail are required for localization and function of chimeric myosin-I at endocytic sites. Overall, this study provides insights into the role of individual myosin-I domains, expands the utility of fission yeast as a simple model system to study the effects of disease-associated *MYO1E* mutations, and supports a model of co-evolution between a myosin motor and its actin track.

## INTRODUCTION

Class I myosins are actin-binding motor proteins that typically function at sites of actin-based membrane deformation during cell shape changes. Humans have 8 class I myosins (Myo1a-h), whose expression and function differs across tissues (1). All class I myosins contain an N-terminal motor domain, a neck region that binds calmodulin or calmodulin-like light chains, and a short membrane-binding tail. While these domains are shared by all members of the myosin-I family, two myosins, Myo1e and Myo1f, known as “long-tailed”, contain an additional proline-rich TH2 domain and an SH3 domain at the C-terminus. “Long-tailed” class I myosins of similar structure also exist in lower eukaryotes, such as yeast, *Dictyostelium,* and *Acanthamoeba,* and are proposed to be the evolutionary ancestor of all common myosins (2). In both unicellular and multicellular organisms, long-tailed class I myosins seem to function in similar physiological processes, including clathrin-mediated endocytosis, pinocytosis, and phagocytosis (3–10). Moreover, these myosins have similar binding partners and even appear at the same stage during endocytosis in diverse organisms. For example, at sites of clathrin-mediated endocytosis in both cultured animal cells and yeast, long-tailed class I myosins function within branched actin networks, assembled by the Arp2/3 complex, where they interact with both membrane phospholipids and members of the WASp family of Arp2/3 complex activators (7,11–16). Class I myosins are recruited to endocytic sites just ahead of actin assembly and are thought to generate force for membrane invagination and scission in both vertebrates and yeast (13,15,17–20).

The conservation of the overall body plan, binding partners, and functional activities between yeast and vertebrate long-tailed class I myosins suggests that the functions of individual protein domains may also be conserved. Given these similarities, we have previously used fission yeast *Schizosaccharomyces pombe*, which has only a single class I myosin Myo1 (7,8), to test the effects of point mutations in the Myo1 motor domain that are homologous to mutations found in the human Myo1e (21). Patients homozygous for these *MYO1E* mutations develop kidney disease (22,23). Introducing these point mutations into yeast Myo1 replicated a loss of myosin-I function phenotype characterized by a severe endocytosis defect resulting in reduced cell growth. Thus, mutations in conserved motor domain residues appear to have similar effects in human and yeast myosin-Is.

Despite structural and functional similarities, the fission yeast Myo1 and human Myo1e also differ in several respects. Myo1 contains an additional region at the C-terminus of the tail, called the central-acidic domain, which can directly bind the Arp2/3 complex to initiate branched actin nucleation (7,11–14). While Myo1e contains a single light-chain binding motif that binds calmodulin (24), *S. pombe* Myo1 contains two IQ motifs, which bind two distinct light-chain proteins, calmodulin Cam1 and calmodulin-like Cam2 (8,25). Lastly, the motor domains of yeast and mammalian class I myosins differ at the TEDS site, a conserved amino acid residue located on a surface loop within the actin-binding site (26). While mammalian myosin-Is have an acidic residue at this site, protozoan and fungal myosin-Is have a phosphorylatable threonine or serine residue at this location (27). Phosphorylation of this threonine or serine by the PAK/Ste20 family kinases regulates myosin ATPase activity and function in amoeba and yeast (11,14,27–33). Together, these differences suggest distinct modes of myosin-I regulation in yeast and mammalian cells.

Although yeast and human class I myosins perform similar functions and have similar protein structure, it is unknown whether these proteins or their individual protein domains are functionally interchangeable. Successful replacement of yeast Myo1 domains with respective domains from human Myo1e would provide insights into the role of individual domains, support future studies introducing protein domains with different biophysical properties to further dissect myosin-I function, and expand the utility of the yeast system for testing the effects of disease-associated mutations. In this study, we tested whether human Myo1e could replace Myo1 in *S. pombe* and found that the human myosin was unable to substitute for yeast Myo1 due to the inability of the Myo1e motor domain to function in yeast cells. However, we found that the Myo1e tail can successfully substitute for the Myo1 tail to support localization and function of myosin-I at endocytic sites, although contributions of individual tail domains differ between these two myosins. Altogether, this work provides novel insights into the roles of individual protein domains and demonstrates that fission yeast can be used as a model system to study the effects of disease-associated Myo1e mutations located in the tail (21).

## RESULTS

### Human Myo1e cannot replace Myo1 in S. pombe

In order to test if human Myo1e, hereafter referred to as HsMyo1e, could functionally substitute for *S. pombe* Myo1, now abbreviated SpMyo1, we replaced the coding sequence of the wild type yeast *myo1^+^* with the sequence coding HsMyo1e at the endogenous *myo1* locus in the yeast genome under control of the endogenous *Pmyo1* promoter. These *myo1Δ::HsMYO1E* cells, like the *myo1Δ* cells (7,8), were viable on regular YES medium at 25°C, but unable to grow in the presence of 1M KCl (Fig. 1A and Table 1). Mutating the TEDS site in the motor domain of HsMyo1e, from glutamate to the phosphorylatable serine, did not improve its function in yeast and *myo1Δ::HsMYO1E(E337S)* cells exhibited the same sensitivity to high salt as *myo1Δ ::HsMYO1E* or *myo1Δ* cells (Fig. 1A and Table 1).

**Figure 1:**
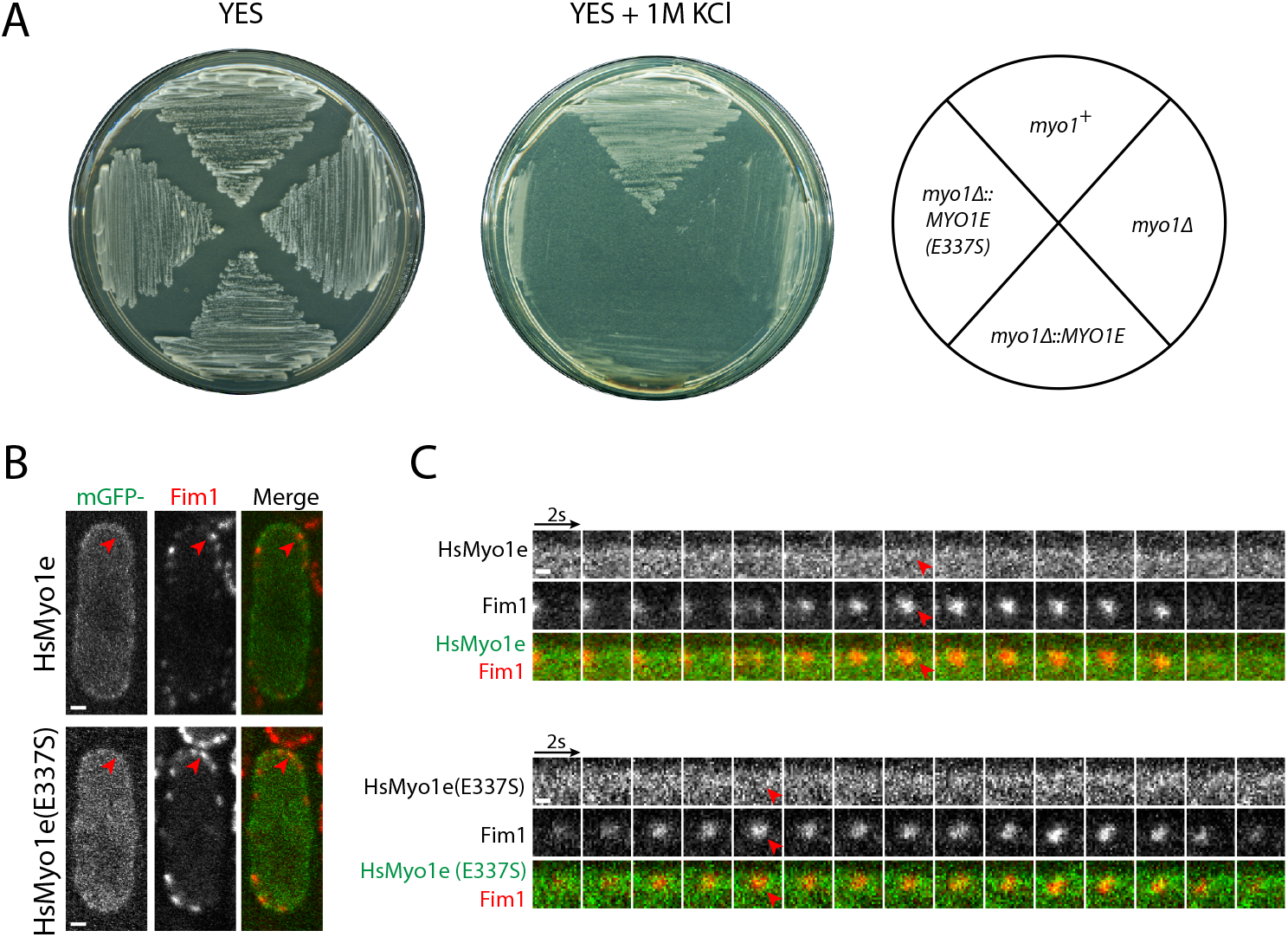
HsMyo1e cannot replace SpMyo1 and fails to localize to sites of endocytosis in *S. pombe.* (A) Analysis of the salt sensitivity of the wild type *(myo1^+^), myo1Δ, myo1Δ::HsMYO1E,* and *myo1Δ::HsMYO1E(E337S) S. pombe* cells. Cells were streaked onto YES agar plates with and without 1M KCl and incubated for several days at 25°C. (B, C) Colocalization analysis of mGFP-tagged (green) HsMyo1e or HsMyo1e(E337S) expressed off of a plasmid under control of *3xPnmt1* promoter in the absence of thiamine with Fim1-mCherry labeled (red) actin patches in *myo1Δ* cells. (B) Single confocal sections through the middle of the cells. Scale bars, 1 μm. (C) Montages of individual patches at 2-second intervals. Scale bars, 0.5 μm. Red arrowheads indicate lack of localization of HsMyo1e or TEDS site mutant HsMyo1e(E337S) with Fim1-mCherry in actin patches.

**Table 1.** Summary of salt sensitivity, localization and actin patch dynamics analyses of HsMyo1e, SpMyo1, and human-yeast myosin-I chimeras in *myo1Δ* cells.

| Name | Description | Promoter | Growth<br>on 1M<br>KCl | Localizes<br>to Actin<br>Patches | Actin patch<br>Internalization<sup>a</sup><br>(%) | Actin patch<br>Lifetime<sup>a</sup><br>(sec) | Colocalizes<br>w/ Cam1 | Colocalizes<br>w/ Cam2 |
| --- | --- | --- | --- | --- | --- | --- | --- | --- |
| CL SpMyo1<sup>b</sup> | *S. pombe* Myo1 | *myo1*<sup>+</sup> | Y | Y | 92 ± 6.6 | 11.6 ± 1.3 | Y | Y |
| HsMyo1e | *H. sapiens* myosin 1e | *myo1*<sup>+</sup> | N | N/A<sup>c</sup> | N.D. | N.D. | N/A<sup>c</sup> | N/A<sup>c</sup> |
| HsMyo1e<br>(E337S) | TEDS site mutant<br>*H. sapiens* myosin 1e | *myo1*<sup>+</sup> | N | N/A<sup>c</sup> | N.D. | N.D. | N/A<sup>c</sup> | N/A<sup>c</sup> |
| HsMyo1e | *H. sapiens* myosin 1e | *nmt1* | N | N | N.D. | N.D. | N | N |
| HsMyo1e<br>(E337S) | TEDS site mutant<br>*H. sapiens* myosin 1e | *nmt1* | N | N | N.D. | N.D. | N | N |
| HsMyo1e tail | HsMyo1e tail | *nmt1* | N | N | N.D. | N.D. | N | N.D. |
| SpMyo1 | *S. pombe* Myo1 | *nmt1* | Y | Y | 96 ± 3.3 | 13.4 ± 2.1 | Y | Y |
| Chimera 1 | HsMyo1e motor-IQ<br>SpMyo1 tail | *nmt1* | N | N | 50 ± 14 | 20 ± 4.4 | N | N |
| Chimera 2 | HsMyo1e motor<br>SpMyo1 IQ1-IQ2-tail | *nmt1* | N | N | 43 ± 7.7 | 20.2 ± 1.7 | N | N |
| Chimera 3 | SpMyo1 motor-IQ1-IQ2<br>HsMyo1e tail | *nmt1* | Y | Y | 77 ± 7 | 14.8 ± 2.5 | Y | Y |
| Chimera 4 | SpMyo1 motor<br>HsMyo1e IQ-tail | *nmt1* | Y | Y | 72 ± 14 | 13.8 ± 2.3 | Y | N |
| Chimera 5 | HsMyo1e motor-IQ<br>SpMyo1 IQ2-tail | *nmt1* | N | N | 32 ± 3.3 | 20.8 ± 1.6 | N | N |
| Chimera 6 | SpMyo1 motor-IQ1<br>HsMyo1e tail | *nmt1* | Y | Y | 80 ± 9 | 12.2 ± 0.6 | Y | N |
| mGFP | Empty vector | *nmt1* | N | N | 46 ± 1.6 | 20.6 ± 3.4 | N | N |

<sup>a</sup>Percent of internalization and lifetimes of actin patches were measured with Fim1-mCherry. Data are presented as ± SD.
<sup>b</sup>CL SpMyo1 designates mGFP-Myo1 expressed from the native *myo1* chromosomal locus under control of endogenous *Pmyo1* promoter.
<sup>c</sup>Data are not available because of the poor expression of Myo1e and Myo1e(E337S) from the *myo1* locus.

Given the differences in codon preferences between yeast and humans (34), we hypothesized that the failure of HsMyo1e to replace SpMyo1 may be due to reduced protein expression in yeast cells. To estimate protein levels, we tagged HsMyo1e and HsMyo1e(E337S) expressed from the *myo1* locus with monomeric GFP (mGFP) at the C-terminus. By fluorescence microscopy, we detected little to no signal for HsMyo1e-mGFP and HsMyo1e(E337S)-mGFP, compared to the whole cell intensity of wild-type SpMyo1, similarly tagged with mGFP and expressed from the native *myo1* locus (Fig. S1). Attempts to detect the mGFP-tagged HsMyo1e and HsMyo1e(E337S) in yeast lysates by western blot were also unsuccessful. The inability of yeast cells to express HsMyo1e from the endogenous *myo1* locus at detectable levels impeded testing whether HsMyo1e can replace SpMyo1. To circumvent this issue, we expressed mGFP-tagged HsMyo1e off of a multi-copy plasmid under control of the full strength, thiamine-repressible *3xPnmt1* promoter. This yeast plasmid contains an autonomously replicating sequence that propagates at a different rate than the yeast genome resulting in considerable variability of the plasmid copy number and consequently variability in protein expression levels amongst cells. To assess the ability of HsMyo1e to function in yeast, we transformed the mGFP-HsMyo1e construct into *myo1Δ* cells expressing fimbrin Fim1 tagged with mCherry as a marker for actin patches. In the absence of thiamine, mGFP-HsMyo1e appeared mostly cytosolic and at elevated levels of expression formed protein aggregates on the membrane (Fig. 1B). We found that mGFP-tagged HsMyo1e and HsMyo1e(E337S) did not localize to sites of endocytosis marked by Fim1-mCherry (Fig. 1C and Table 1). Even upon overexpression, HsMyo1e failed to rescue the inability of *myo1Δ* cells to grow on high salt (Fig. 2 and Table 1). Thus, HsMyo1e is not able to substitute for SpMyo1 in fission yeast cells, even when expressed at levels greater than the endogenous SpMyo1.

**Figure 2:**
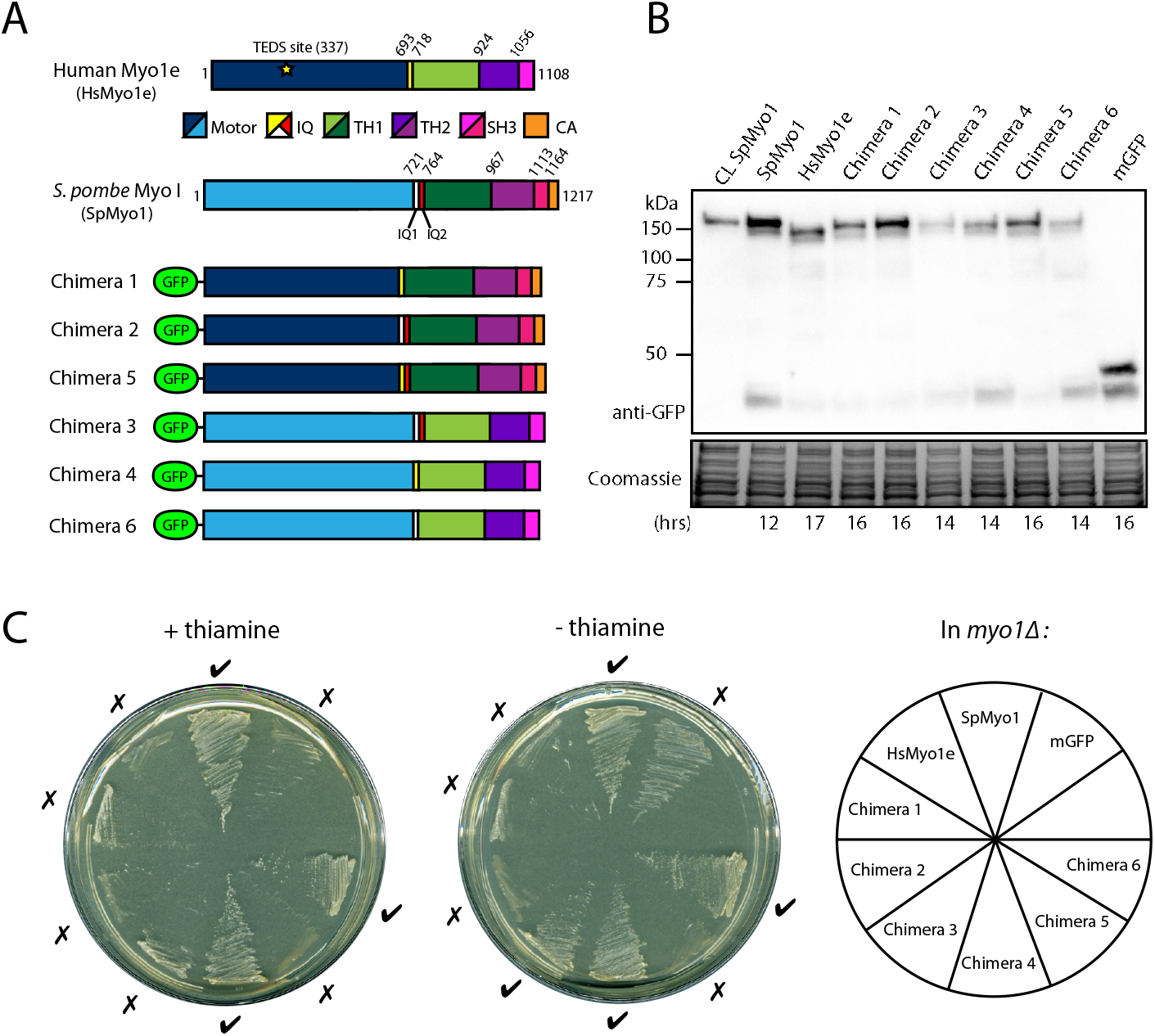
Human-yeast myosin-I chimeras require the SpMyo1 motor domain to rescue *myo1Δ.* (A) Domain maps of *H. sapiens* Myo1e (HsMyo1e), *S. pombe* Myo1 (SpMyo1), and six human-yeast myosin-I chimeric constructs tagged with mGFP at the N-terminus. Chimeras 1, 2, and 5 contain the HsMyo1e motor domain and the SpMyo1 tail. Chimeras 3, 4, and 6 contain the SpMyo1 motor domain and the HsMyo1e tail. IQ, light chain binding IQ motif; TH, tail homology domain; SH3, Src Homology 3 domain; CA, central-acidic domain. (B) Western blot analysis of total protein lysates of cells expressing mGFP-Myo1 (CL SpMyo1) from native *myo1* locus under control of endogenous *Pmyo1* promoter or *myo1Δ* cells expressing mGFP-tagged SpMyo1, HsMyo1e, human-yeast myosin-I chimeras, or mGFP alone from plasmids under control of *3xPnmt1* promoter. The times cells were grown in the absence of thiamine to induce protein expression are indicated at the bottom. Blots were probed with anti-GFP antibody. Coomassie stained gel serves as a loading control. (C) Analysis of salt sensitivity of *myo1Δ* cells expressing mGFP alone, mGFP-tagged SpMyo1, HsMyo1e, or SpMyo1-HsMyo1e chimeras off of plasmid under control of *3xPnmt1* promoter in the presence and the absence of thiamine at 25°C on EMM agar plates containing 1M KCl. These cells also express actin patch marker Fim1-mCherry. Growth (✓) or lack of growth (×) is indicated around the plates.

### Human-yeast myosin-I chimeras require the SpMyo1 motor domain to rescue myo1Δ

Given the inability of HsMyo1e to replace SpMyo1, we next generated six chimeric constructs composed of varying combinations of HsMyo1e and SpMyo1 domains to determine which protein domain makes HsMyo1e unable to function in yeast (Fig. 2A). We defined the boundaries of HsMyo1e and SpMyo1 protein domains based on the alignment of their amino acid sequences (Fig. S2) and previous reports (7,35). Three of the chimeras, Chimeras 1, 2, and 5, contained the HsMyo1e motor domain paired with the SpMyo1 tail, and three others, Chimeras 3, 4, and 6, contained the SpMyo1 motor domain and the HsMyo1e tail. To ascertain the functional importance of the IQ domains, we shuffled the IQ domains from SpMyo1 and HsMyo1e in different combinations among the chimeras. All chimeras were tagged at the N-terminus with mGFP and expressed off of a multi-copy plasmid under control of the thiamine-repressible *3xPnmt1* promoter. The mGFP-tagged myosin chimeras were then transformed into a *myo1Δ* strain expressing Fim1-mCherry as a marker of actin patches and assessed for protein expression and the ability to support growth of *myo1Δ* cells on high salt (Fig. 2 and Table 1). We used mGFP-tagged SpMyo1 and mGFP alone expressed from the same multi-copy plasmids as positive and negative controls, respectively. Western blot analysis revealed that all chimeric myosins were expressed as full-length proteins of expected molecular weights (Fig. 2B). Interestingly, HsMyo1e and the chimeras containing the HsMyo1e motor domain took longer to express than SpMyo1 and chimeras containing the SpMyo1 motor domain (Fig. 2B). Growth assays in the presence of 1M KCl revealed that the chimeras containing the SpMyo1 motor domain with the HsMyo1e tail partially rescued the growth defect of *myo1Δ* cells on high salt, similar to the SpMyo1 positive control but to a lesser degree (Fig. 2C). Curiously, these constructs were able to support growth on high salt even when expressed at low levels in the presence of thiamine. In contrast, the chimeras with the HsMyo1e motor domain, like HsMyo1e and the mGFP control, completely failed to rescue the growth defect of *myo1Δ* cells on high salt (Fig. 2C). These results suggest that the HsMyo1e motor domain is responsible for the inability of HsMyo1e to replace SpMyo1 and that the function of SpMyo1 requires the endogenous motor domain.

### Human-yeast myosin-I chimeras require the SpMyo1 motor domain for localization to actin patches

To determine if the chimeras’ ability to rescue *myo1Δ* growth defects correlated with their ability to localize to actin patches, we examined their localization in *myo1Δ* cells expressing Fim1-mCherry (Fig. 3 and Table 1). Fluorescence microscopy revealed that indeed the rescuing Chimeras 3, 4, and 6, composed of the SpMyo1 motor domain and HsMyo1e tail, localized to dynamic actin patches marked by Fim1-mCherry (Fig. 3). However, unlike SpMyo1, Chimeras 3, 4, and 6 also strongly outlined the entire cell cortex. These chimeras are likely targeted to the cell cortex by Myo1e tail since mGFP-tagged HsMyo1e tail alone similarly outlined the cell cortex but failed to localize to actin patches (Fig. S3). Since the SpMyo1 motor domain alone also fails to localize to actin patches (7), both the SpMyo1 motor domain and HsMyo1e tail are needed for these chimeras to localize to actin patches.

**Figure 3:**
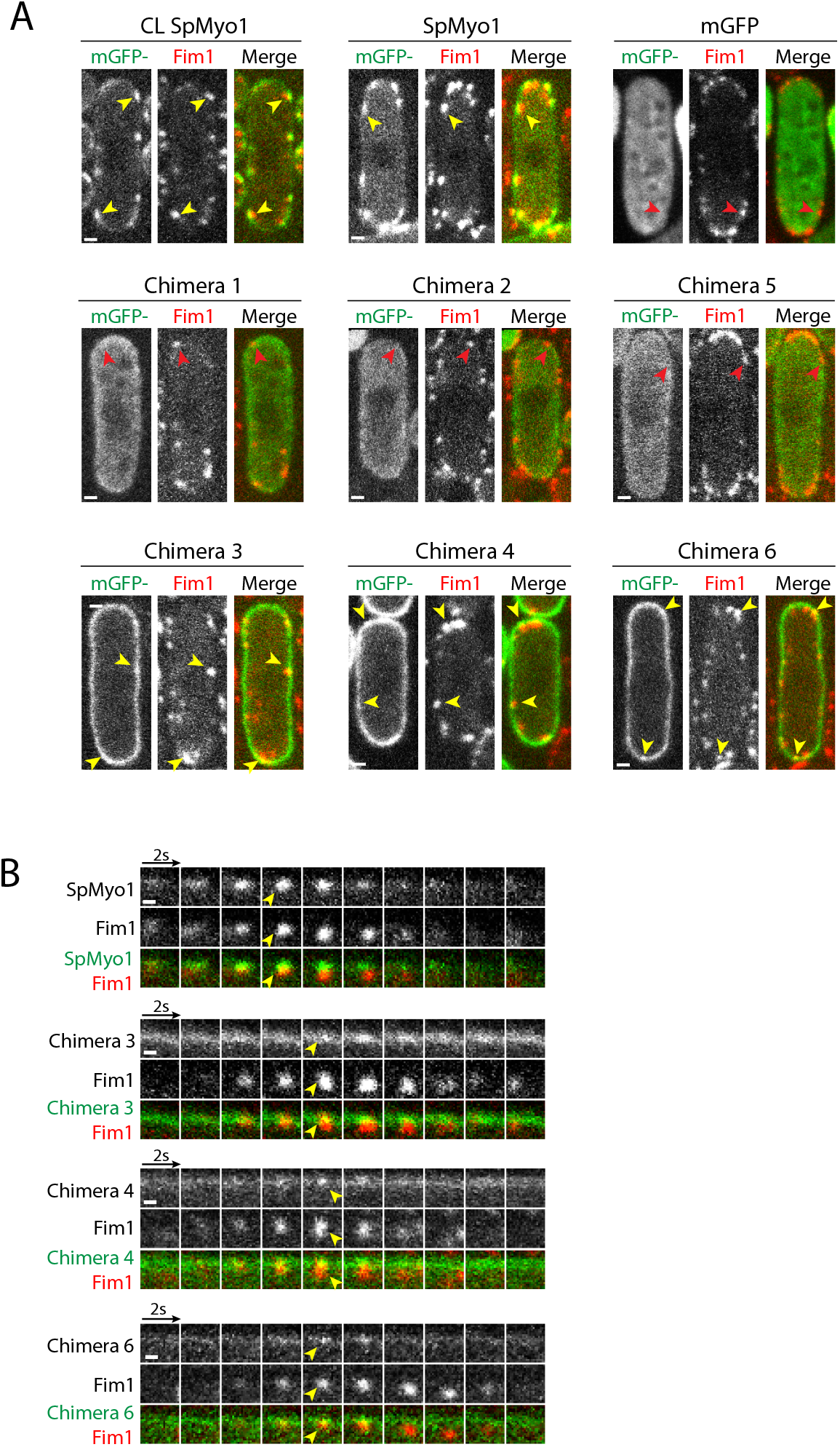
Human-yeast myosin-I chimeras require the SpMyo1 motor domain for localization to actin patches. (A, B) Analysis of colocalization of mGFP-tagged (green) SpMyo1 or SpMyo1-HsMyo1e chimeras with Fim1-mCherry (red) in actin patches in *myo1Δ* cells. In control, mGFP-Myo1 (CL SpMyo1) was expressed from the native *myo1* locus under control of the endogenous *Pmyo1* promoter in wild-type cells expressing Fim1-mCherry. mGFP-tagged SpMyo1, mGFP alone, and mGFP-tagged human-yeast myosin-I chimeras were expressed from plasmids under control of *3xPnmt1* promoter for 12-18 hours in the absence of thiamine in *myo1Δ* cells expressing Fim1-mCherry. (A) Representative images in single confocal sections through the middle of the cells. Scale bars, 1 μm. (B) Montages of individual patches at 2-second intervals. Scale bar, 0.5 μm. Yellow and red arrowheads indicate the presence and the absence of myosin in Fim1-mCherry-labeled actin patches, respectively.

Examination of Chimeras 1, 2, and 5, which failed to rescue growth of *myo1Δ* cells under high salt conditions revealed that these chimeras, composed of the HsMyo1e motor domain and SpMyo1 tail, failed to localize to actin patches marked by Fim1-mCherry (Fig. 3 and Fig. S3). All three chimeras exhibited mostly cytosolic localization and only mild accumulation at the cell cortex, compared to Chimeras 3, 4, and 6. While Chimera 1 lacking the IQ2 motif of SpMyo1 lightly outlined the entire cortex, Chimeras 2 and 5, which both contain the IQ2 motif of SpMyo1 in addition to the SpMyo1 tail, localized to cortical thread-like structures reminiscent of eisosomes (36) in about 35-40% of the cells (Fig. S3B). To verify this, we transformed mGFP-tagged Chimeras 2 and 5 into *myo1Δ* cells expressing a known eisosome marker Fhn1 tagged with mCherry (36). However, we could not detect convincing colocalization of the two proteins, suggesting that the cortical linear structures recruiting Chimeras 2 and 5 may not be eisosomes, or that tagging Fhn1 with mCherry interferes with the localization of these chimeras to eisosomes. Overall, this analysis revealed that the HsMyo1e motor domain combined with the SpMyo1 tail is not sufficient for localizing chimeric constructs to actin patches.

### SpMyo1 motor - HsMyo1e tail chimeras support dynamics and internalization of actin patches

Since chimeras containing the SpMyo1 motor domain were capable of localizing to actin patches and rescuing the salt-sensitive growth defect of *myo1Δ* cells, we examined the effect of human-yeast myosin-I chimeras on actin patch dynamics and endocytic internalization in *myo1Δ* cells expressing Fim1-mCherry (Fig. 4 and Table 1). We measured the time courses of intensities and distance traveled for mGFP-tagged myosin and Fim1-mCherry in individual actin patches in live cells and quantified the percent of internalizing actin patches (Fig. 4 and Fig. S4). As not all patches in chimera expressing cells internalized, we generated average time courses (Fig. 4A and Fig. S4A) by aligning individual time courses to the time Fim1-mCherry reached peak intensity (time zero), which in wild-type cells corresponds to the initiation of endocytic internalization (37). Because the plasmid-based exogenous expression was highly variable from cell to cell, we compared the expression level of all constructs in individual cells to mGFP-tagged SpMyo1 expressed from the endogenous locus by measuring total cell intensities. Using the SpMyo1 control expressed off the plasmid, we determined whether varying levels of SpMyo1 expression have any impact on accumulation of SpMyo1 in actin patches and actin patch dynamics. This analysis revealed that SpMyo1 expression level correlated weakly with peak accumulation of SpMyo1 in patches, so that up to 2-fold more SpMyo1 accumulated in patches in cells expressing 1.5-3.5-fold more SpMyo1 than the endogenous level (Fig. S4B). This increased patch accumulation of SpMyo1 in higher expressing cells had, however, no effect on peak accumulation of Fim1-mCherry (Fig. S4C and D). On the other hand, in the complete absence of SpMyo1, peak accumulation of Fim1-mCherry was reduced by ~40% (Fig. 4B). To avoid potential artifacts due to variable expression levels, we chose to track actin patch dynamics only in cells that expressed myosin chimeras at the levels within the range of two-fold higher or lower than the endogenous level of SpMyo1. Expression of the chimeras in *myo1^+^* cells resulted in ~100% patch internalization rate, indicating that none of the myosin constructs exhibited dominant negative effects (Fig. S4E).

**Figure 4:**
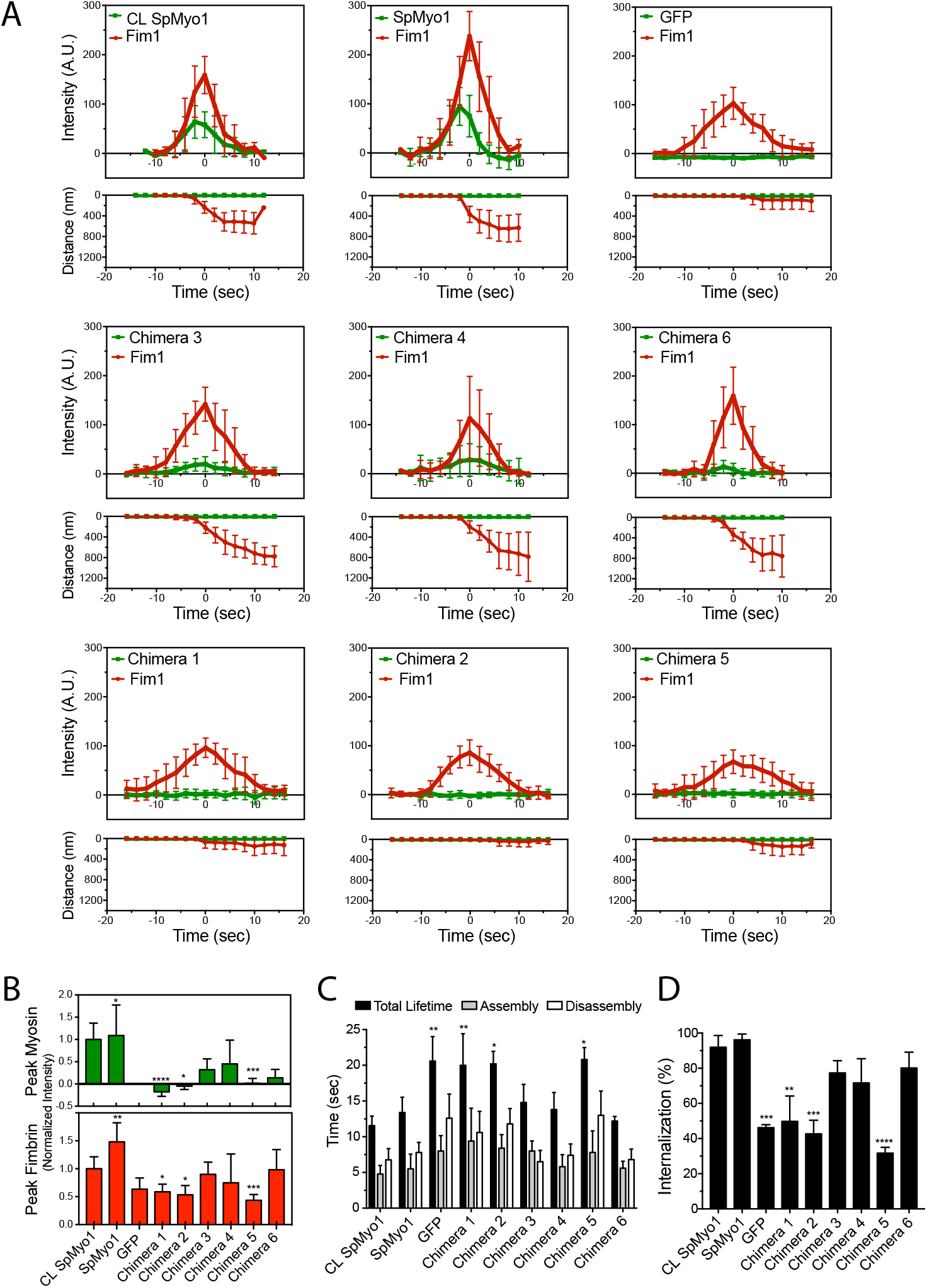
SpMyo1 motor - HsMyo1e tail chimeras partially rescue dynamics of endocytic internalization. (A-D) Analysis of localization and dynamics of mGFP-tagged human-yeast myosin-I chimeras and Fim1-mCherry in actin patches in *myo1Δ* cells. mGFP-SpMyo1, mGFP alone, and mGFP-tagged chimeras were expressed from plasmids under control of *3xPnmt1* promoter for 12-18 hours in the absence of thiamine in *myo1Δ* cells expressing Fim1-mCherry. In control (CL SpMyo1), the dynamics of mGFP-Myo1 expressed from endogenous *myo1* locus and Fim1-mCherry were tracked in wild-type cells. (A) Time courses of (upper panels) average fluorescence intensity (± SD) and (lower panels) average distance traveled (± SD) of mGFP-tagged (green squares) SpMyo1 or chimeras and Fim1-mCherry (red circles) in actin patches in *myo1Δ* cells or, in control, wild-type cells. The time courses of cortical background-subtracted intensities and distances from the origin for individual patches were aligned to the peak of Fim1-mCherry patch intensity (time zero) and averaged at each time point. (B) Bar graphs (mean ± SD) of peak intensities from averaged plots of mGFP-tagged myosin (green) and Fim1-mCherry (red) in actin patches in *myo1Δ* cells normalized to intensities in CL SpMyo1 cells. (C) Bar graph (mean ± SD) of assembly time (grey), disassembly time (white), and total lifetime (black) of Fim1-mCherry in actin patches in WT cells or *myo1Δ* cells expressing mGFP-tagged SpMyo1 or SpMyo1-HsMyo1e chimeras. (D) Bar graph of percent internalization (± SD) of actin patches marked with Fim1-mCherry in WT cells or in *myo1Δ* cells expressing mGFP-tagged SpMyo1 or human-yeast myosin-I chimeras. N=9-12 patches in 3-6 cells in (A-C) and N=23-43 patches in 3-6 cells in (D). Asterisks indicate a significant difference from WT control as determined by a one-way ANOVA, * p<0.05, ** p<0.01, ***p<0.001, ****p<0.0001.

In agreement with the ability to localize to actin patches and support growth in the presence of high salt, Chimeras 3, 4, and 6 partially rescued the actin patch dynamics and endocytic internalization defects observed in *myo1Δ* cells. In *myo1Δ* cells expressing mGFP alone, the times of assembly, disassembly and total lifetime of Fim1-mCherry in patches are increased and 40-60% of patches fail to internalize (Fig. 4). Expression of SpMyo1 off of the plasmid completely restored actin patch dynamics to wild-type levels: Fim1-mCherry lifetime in patches was 13.4±2.1 seconds, similar to 11.5±1.3 seconds observed in cells expressing SpMyo1 from the endogenous locus, and 90-100% of patches internalized (Fig. 4). Chimeras 3, 4, and 6 accumulated in patches at 20%, 30%, and 10% levels of endogenously expressed SpMyo1 (Fig. 4A and 4B) and partially rescued actin patch dynamics (Fig. 4C and 4D). However, these accumulation levels may be underestimated due to the enrichment of these myosins at the cell cortex making measurements more challenging. Regardless, in cells expressing these chimeras, peak accumulation and lifetimes of Fim1-mCherry in patches were similar to those observed in cells expressing SpMyo1 (Fig. 4C) and the percent of successfully internalized patches increased to 70-80% (Fig. 4D). Thus, the combination of the SpMyo1 motor domain with the HsMyo1e tail is able to support endocytosis in fission yeast.

In contrast, Chimeras 1, 2, and 5 that failed to localize to patches and rescue *myo1Δ* growth under high salt conditions did not rescue actin patch dynamics and internalization defects in *myo1Δ* cells (Fig. 4 and Table 1). Cells expressing these chimeras exhibited significantly longer Fim1-mCherry patch lifetimes (Chimera 1: 20±3.4s; Chimera 2: 20.2±1.7s; Chimera 5:20.8±1.6s) and reduced accumulation of Fim1-mCherry in patches (Fig. 4A-C and Table 1). Furthermore, patch internalization in *myo1Δ* cells expressing these chimeras remained at 30–50%, similar to *myo1Δ* cells (Fig 4D and Table 1). Thus, the SpMyo1 tail cannot support endocytic function when paired with the HsMyo1e motor.

### HsMyo1e IQ motif recruits Cam1 but not Cam2, in S. pombe

To further dissect the functionality of these human-yeast myosin-I chimeras, we tested whether they could bind and recruit myosin light chains, which regulate myosin activity. Vertebrate Myo1e has a single IQ motif bound by calmodulin that inhibits myosin ATPase activity in the presence of Ca^2^+^^ (24). *S. pombe* Myo1 has two IQ motifs, IQ1 bound by calmodulin Cam1 and IQ2 bound by the calmodulin-like light chain Cam2 (8,25). The IQ motif of HsMyo1e shares greater amino acid sequence similarity with the first IQ motif of SpMyo1 (38%) than with the second IQ motif (23%) (Fig. S2). Thus, we hypothesized that yeast Cam1 may interact with the HsMyo1e IQ domain. To test if the HsMyo1e IQ motif can recruit Cam1, we transformed the mGFP-tagged chimeric constructs into a *myo1Δ* strain expressing Cam1 tagged with mCherry (Fig. 5, Fig. S5, and Table 1). As expected, the mGFP-SpMyo1 control and mCherry-Cam1 colocalized in 100% of endocytic patches (Fig. 5 and Fig. S5A). The SpMyo1 motor – HsMyo1e tail Chimeras 3, 4, and 6, previously observed to localize to endocytic actin patches, also showed colocalization with Cam1 in dynamic cortical patches (Fig. 5A and B), regardless of whether they had the SpMyo1 IQ1 motif (Chimeras 3 and 6) or HsMyo1 IQ motif (Chimera 4). On average, Chimeras 3, 4 and 6 colocalized with mCherry-Cam1 in ~73% of actin patches (Fig. S5A), likely an underestimate due to reduced levels of these chimeras in patches and abundance of Cam1 in the cytoplasm. At high levels of expression, these three chimeras strongly outlined the cortex and recruited a significant amount of mCherry-Cam1 from the cytoplasm to the cortex (Fig. S5B). This recruitment was specific for the IQ motif-Cam1 interaction as cells overexpressing mGFP-HsMyo1e tail produced no such effect. The ability of HsMyo1e IQ motif to recruit Cam1 was also apparent in Chimera 1, which colocalized with Cam1 over the entire cortex. Chimeras 2 and 5, which contain the SpMyo1 IQ1 motif and HsMyo1e IQ motif, respectively, were observed recruiting Cam1 to the cortical thread-like structures (Fig. S6A). Thus, the HsMyo1e IQ domain appears to be capable of binding Cam1 and recruiting it to actin patches and other cortical structures.

**Figure 5:**
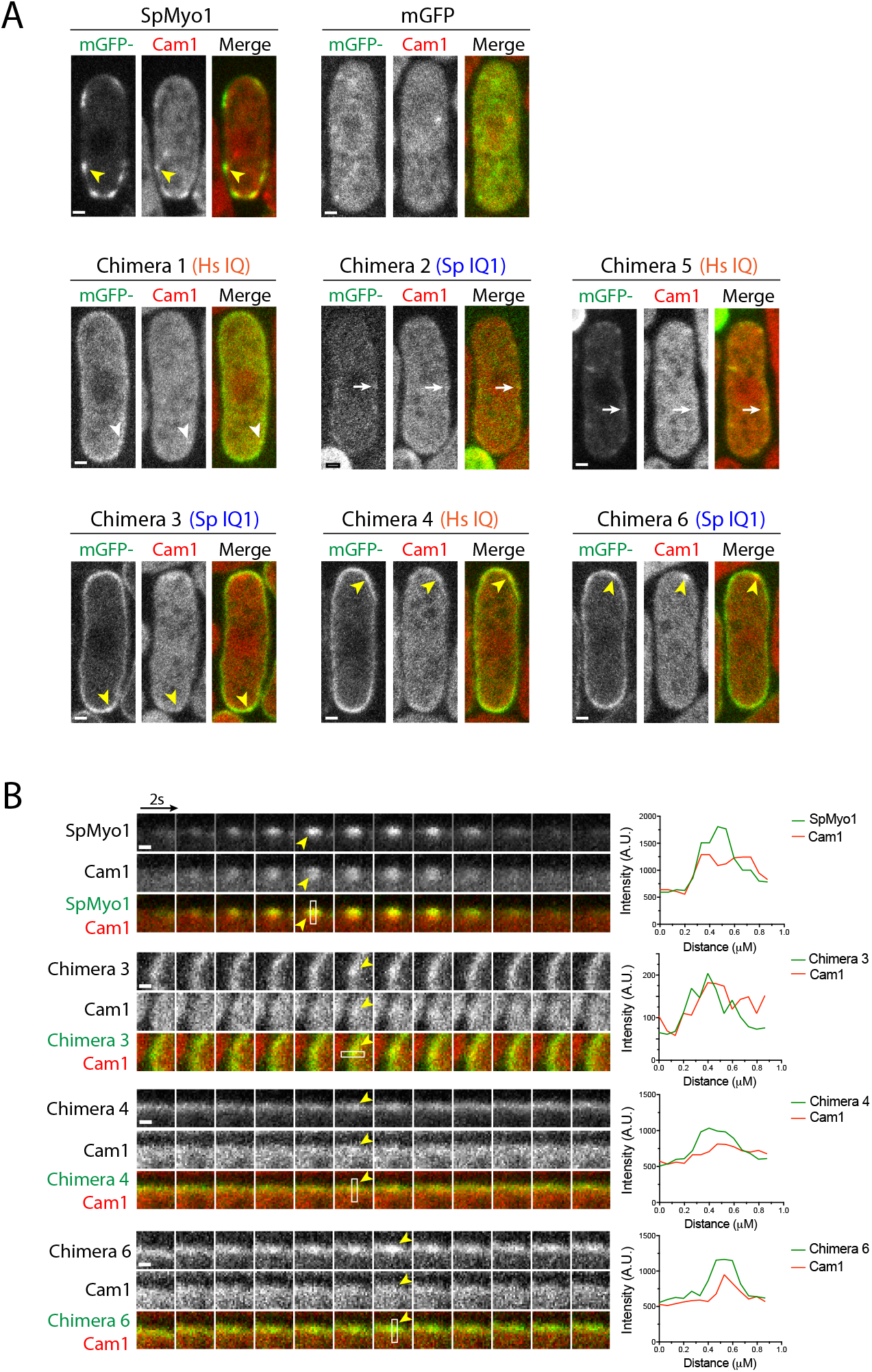
HsMyo1e IQ and SpMyo1 IQ1 motifs recruit calmodulin Cam1 in *S. pombe.* (A-C) Colocalization analysis of mGFP-tagged (green) human-yeast myosin-I chimeras and mCherry-tagged calmodulin Cam1 (red) in *myo1Δ* cells. mGFP alone, mGFP-tagged SpMyo1 and chimeras were expressed from plasmids under control of *3xPnmt1* promoter for 12-18 hours in the absence of thiamine. (A) Single confocal sections through the middle of the cells. The orange Hs IQ and blue Sp IQ1 labels indicate the presence of the HsMyo1e IQ motif and the SpMyo1 IQ1 motif, respectively. Scale bars, 1 μm. (B) Montages of individual patches at 2-second intervals and line scans of fluorescence intensity in patches. White boxes mark areas used for line scans. Scale bar, 0.5 μm. Yellow arrowheads indicate colocalization of Cam1 with SpMyo1 and Chimeras 3, 4 and 6 in actin patches. White arrowheads indicate colocalization of Cam1 with Chimera 1 at cell cortex and white arrows indicate colocalization of Cam1 with Chimeras 2 and 5 in eisosome-like punctate cortical structures.

Cam2 binding to the lever arm of SpMyo1 has been found to maximize myosin motility *in vitro* (25). In the absence of Cam2, SpMyo1 still localizes to actin patches, but in the absence of SpMyo1 or when Cam2-IQ2 motif binding is otherwise disrupted, Cam2 moves to larger stationary puncta at the poles as well as mobile cytosolic puncta (25). To determine whether the chimeric proteins were capable of recruiting Cam2, we transformed the mGFP-tagged myosin constructs into a *myo1Δ* strain expressing Cam2 tagged with mCherry (Fig. 6, Fig. S6 and Table 1). Upon expression of mGFP-SpMyo1, Cam2 was recruited to dynamic actin patches and completely colocalized with SpMyo1 (Fig. 6). As observed previously, Cam2 in *myo1Δ* cells expressing mGFP alone localized in bright puncta at the poles. Chimeras 1, 4 and 6 lacking the SpMyo1 IQ2 motif failed to recruit Cam2 or to colocalize with Cam2 puncta. The lack of colocalization between Cam2 and Chimeras 1 and 4 containing the HsMyo1e IQ1 motif or Chimera 6 containing the SpMyo1 IQ1 motif highlights the specificity of Cam2 binding to the SpMyo1 IQ2 motif. Chimera 3, the only chimera to both localize to endocytic patches and contain the IQ2 domain, colocalized with Cam2 in dynamic cortical patches (Fig. 6B). Chimeras 2 and 5 that localized to cortical thread-like structures and contain the IQ2 motif recruited Cam2 to these eisosome-like structures, just as they did with Cam1 (Fig. S6B). Thus, Cam2 only bound chimeras that contained the SpMyo1 IQ2 motif.

**Figure 6:**
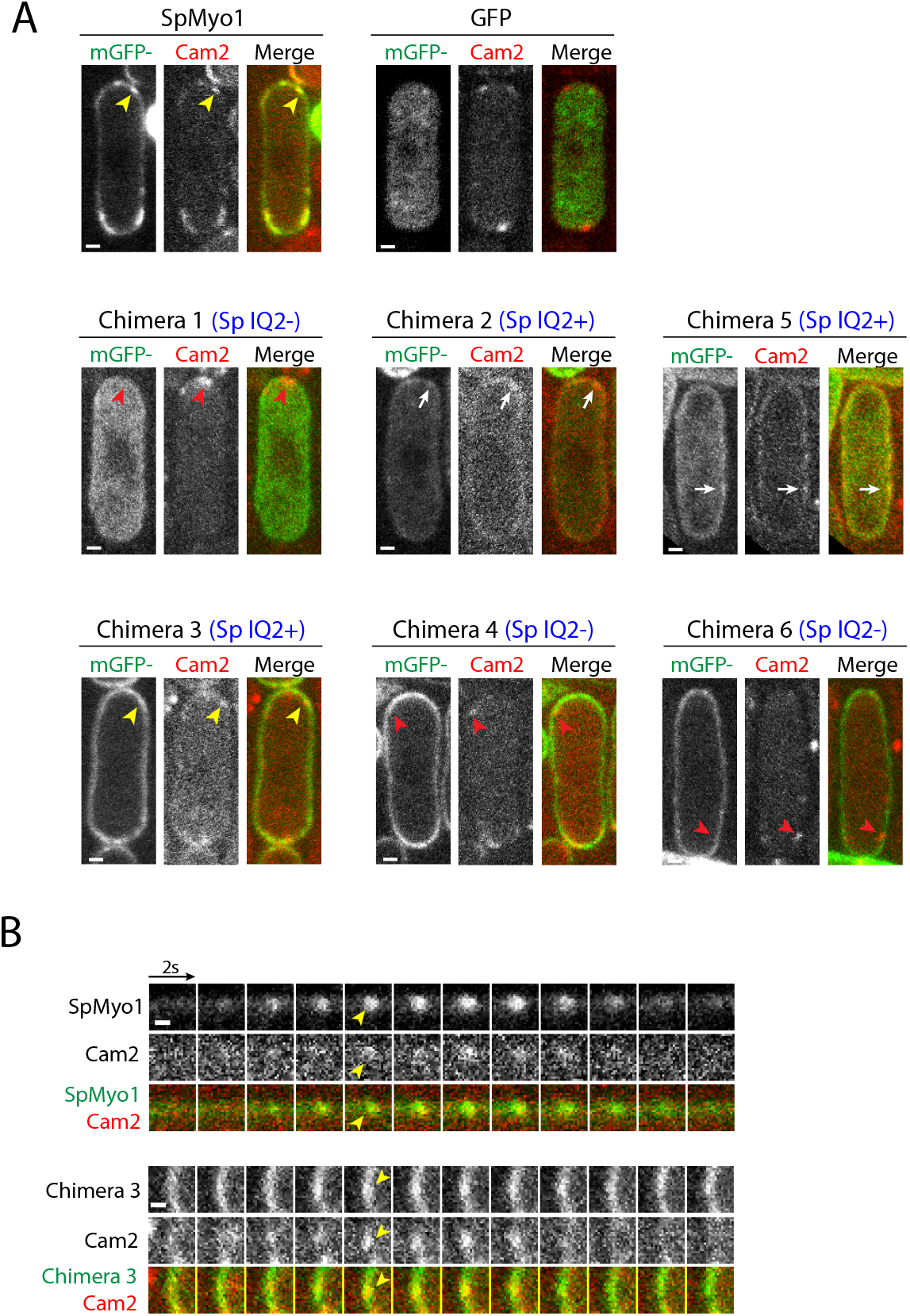
SpMyo1 IQ2 motif recruits calmodulin-related light chain Cam2 in *S. pombe.* (A-C) Colocalization analysis of mGFP-tagged (green) human-yeast myosin-I chimeras and mCherry-tagged calmodulin-like light chain Cam2 (red) in *myo1Δ* cells. mGFP alone, mGFP-tagged SpMyo1 and chimeras were expressed from plasmids under control of *3xPnmt1* promoter for 12-18 hours in the absence of thiamine. (A) Single confocal sections through the middle of the cells. The blue Sp IQ^+^/− labels indicate the presence or the absence of the SpMyo1 IQ2 motif. Scale bars, 1 μm. (B) Montages of individual patches at 2-second intervals. Scale bar, 0.5 μm. Yellow arrowheads indicate colocalization of Cam2 with SpMyo1 and Chimera 3 in actin patches. White arrows indicate colocalization of Cam2 with Chimeras 2 and 5 in eisosome-like punctate cortical structures. Red arrowheads indicate Cam2-only puncta that form in the absence of SpMyo1 IQ2 motif.

### The SpMyo1 motor – HsMyo1e tail chimera requires the TH1 and TH2-SH3 domains for localization and function

Given the ability of the SpMyo1 motor – HsMyo1e tail chimeras to localize to endocytic actin patches and to rescue *myo1Δ* endocytosis defects, we sought to determine which HsMyo1e tail domains are minimally required for their localization and function. Using Chimera 3, we created two tail domain deletion constructs: one lacking the TH1 domain and another lacking the TH2 and SH3 domains (Fig. 7A). For comparison, we made similar deletions in SpMyo1, one removing the membrane-binding TH1 domain and another removing the C-terminal TH2, SH3, and CA domains. These mGFP-tagged constructs were transformed into the *myo1Δ* strain expressing Fim1-mCherry and tested for salt sensitivity, localization to actin patches, and the ability to rescue the *myo1Δ* patch internalization defect. Immunoblotting confirmed that all constructs were expressed at the expected molecular weights (Fig. 7B). Testing for growth in the presence of high salt revealed that both Chimera 3 tail domain deletion mutants failed to rescue the growth of *myo1Δ* cells (Fig. 7C). In line with this finding, Chimera3ΔTH1 and Chimera3ΔTH2-SH3 failed to localize to actin patches marked with Fim1-mCherry and were mostly cytosolic (Fig. 7D and E). The fraction of internalizing patches in cells expressing these constructs remained at ~50%, similar to *myo1Δ* cells (Fig. 7F). Conversely, analogous deletions of tail domains from SpMyo1 resulted in drastically different behaviors. The SpMyo1ΔTH2-SH3-CA rescued the growth of *myo1Δ* cells in the presence of high salt, localized to dynamic actin patches, and partially rescued the patch internalization defect, increasing the fraction of internalizing patches to ~80% (Fig. 7C, D, and F and Fig. S7). In contrast, SpMyo1ΔTH1 failed to rescue salt-sensitive growth (Fig. 7C) and the patch internalization defect in *myo1Δ* cells (Fig. 7F), even though SpMyo1ΔTH1 clearly localized to actin patches marked with Fim1-mCherry (Fig. 7D). Remarkably, unlike wild-type SpMyo1 that remains at the plasma membrane during actin patch internalization, SpMyo1ΔTH1 left the membrane and moved away together with Fim1-mCherry in a fraction of internalizing actin patches (Fig. S7A). This behavior was also observed in the presence of SpMyo1 in wild-type cells (Fig. S7B). This finding emphasizes the importance of the TH1 domain for anchoring SpMyo1 at the membrane so that it can function in endocytic internalization. Intriguingly, while neither TH1 nor TH2-SH3-CA are not required for SpMyo1 localization to endocytic patches, both the TH1 and TH2-SH3 domains in the HsMyo1e tail are needed for localization and function of SpMyo1 motor – HsMyo1e tail chimeras.

**Figure 7:**
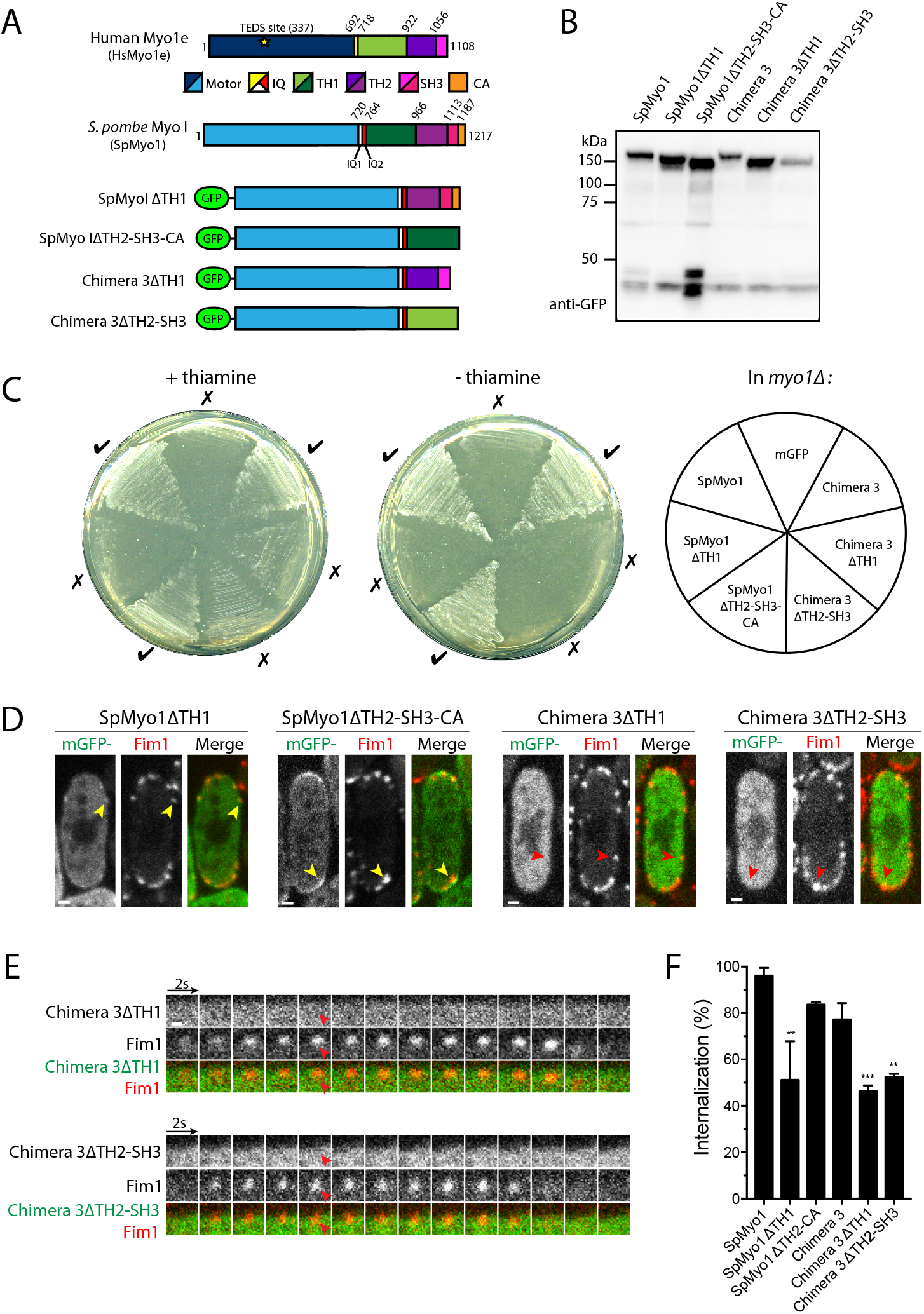
SpMyo1 motor – HsMyo1e tail chimera requires both Myo1e TH1 and TH2-SH3 domains to rescue *myo1Δ* and to localize to actin patches. (A) Domain maps of *H. sapiens* Myo1e (HsMyo1e), *S. pombe* Myo1 (SpMyo1) and mGFP-tagged tail domain deletion constructs of SpMyo1 and SpMyo1-HsMyo1e Chimera 3. (B) Western blot of total protein lysates of *myo1Δ* cells expressing indicated mGFP-tagged myosin constructs. The constructs were expressed from plasmids under control of *3xPnmt1* promoter for 12-15 hours in the absence of thiamine. The blot was probed with an anti-GFP antibody. (C) Analysis of salt sensitivity of *myo1Δ* cells expressing indicated mGFP-tagged myosin constructs from plasmids under control of *3xPnmt1* promoter in the presence or absence of thiamine at 25°C on EMM agar plates containing 1M KCl. The cells also expressed Fim1-mCherry. Growth (✓) or lack of growth (×) is indicated around the plate. (D-E) Colocalization analysis of indicated mGFP-tagged (green) myosin constructs expressed from plasmids under control of *3xPnmt* promoter in the absence of thiamine and Fim1-mCherry (red) in actin patches in *myo1Δ* cells. (D) Single confocal sections through the middle of the cells. Scale bars, 1 μm. (E) Montages of individual patches at 2-second intervals. Scale bar, 0.5 μm. Yellow arrowheads indicate colocalization and red arrowheads indicate lack of colocalization of myosin with Fim1-mCherry in actin patches. (F) Bar graph of percent internalization (± SD) of Fim1-mCherry patches in in *myo1Δ* cells expressing indicated myosin constructs. Data for SpMyo1 and Chimera 3 are copied from Figure 4C. N = 132-223 patches in 11-19 cells. Asterisks indicate a significant difference from the mGFP-tagged SpMyo1 expressed from a plasmid under control of *3xPnmt1* promoter as determined by one-way ANOVA, ** p<0.01, ***p<0.001, ****p<0.0001.

## DISCUSSION

In this study, we set out to determine whether the conservation of overall functions and general protein organization between yeast and human myosin-Is reflects the conserved properties of individual protein domains or whether their protein domain properties have diverged while retaining a common function in endocytosis and overall protein organization. We found that HsMyo1e cannot replace SpMyo1 as a functional motor at sites of endocytosis. By creating human-yeast myosin-I chimeras, we tracked the failure of HsMyo1e to function in yeast to its motor domain as chimeras composed of the HsMyo1e motor and SpMyo1 tail could not replicate myosin-I function. Meanwhile, chimeras composed of the SpMyo1 motor domain and HsMyo1e tail localized to endocytic actin patches and partially rescued the endocytic defects of *myo1Δ* cells. Further dissection revealed that individual tail domains of HsMyo1e and SpMyo1 make differing contributions to myosin-I localization and function. Overall, these findings point to co-evolution of myosin motor with the actin track, illuminate the role of individual protein domains in myosin-I localization and function, and set limits on which protein domains can be swapped with respective domains from humans and other organisms to study myosin-I function and the effects of disease-associated mutations.

### Evolutionary conservation of myosin-I motor domain with its actin track

The inability of HsMyo1e motor domain to work properly with yeast actin appears to be the primary reason for the failure of HsMyo1e and HsMyo1e motor domain containing chimeras in *S. pombe.* Indeed, while HsMyo1e expressed at extremely low levels from the *myo1* locus and HsMyo1e motor domain containing chimeras took longer to express, all chimeric constructs reached expression levels comparable or exceeding the endogenous level of SpMyo1 when expressed off of a plasmid. Moreover, all constructs were expressed as full-length proteins of expected molecular weights and did not aggregate at moderate expression levels. These results provide evidence that poor expression or misfolding were not significant factors contributing to HsMyo1e motor domain failure.

The HsMyo1e motor domain may not function in *S. pombe* due to differences in regulation of myosin-I activity in yeast and mammals, or the absence of specific myosin or actin binding proteins. While phosphorylation at the TEDS site is critical for SpMyo1 localization and function(33), HsMyo1e contains a glutamate residue at the TEDS site and therefore is expected to be constitutively active. The Glu-to-Ser mutation at the TEDS site failed to improve HsMyo1e functionality in yeast, suggesting that the lack of regulation at the TEDS site is not the main factor compromising HsMyo1e function in yeast. Alternatively, yeast kinases may not recognize the phosporylation site on the HsMyo1e motor. The other mechanism of regulation of myosin-I activity is mediated by calmodulin or calmodulin-like myosin light chains that bind the IQ motifs in a myosin neck region, thereby stabilizing the myosin lever arm for efficient force production (38–40). Yet recruitment of Cam1 to actin patches by the HsMyo1e IQ motif suggests that HsMyo1e motor domain failure is not due to the lack of stabilization of the lever arm by calmodulin. Furthermore, earlier work demonstrated that budding yeast and vertebrate calmodulins are functionally interchangeable (41). Among actin binding proteins, tropomyosins are particularly important for regulating myosin activity and also serving as gatekeepers controlling the localization of myosins and other proteins by binding actin filaments (42). However, both mammalian and fungal myosin-Is are excluded from tropomyosin-occupied actin filaments and tropomyosin is not present at actin patches in yeast (43–45).

Therefore, it is more likely that the inability of HsMyo1e motor domain to function in yeast is due to species related differences in the interaction of actin and myosin. *S. pombe* actin significantly differs from mammalian actin kinetically, having a faster nucleotide exchange rate, phosphate dissociation rate, and F-actin nucleation rate by the Arp2/3 complex (46), which indicates that differences in actin properties and actin-myosin interaction could be the cause for HsMyo1e failure to function in *S. pombe.* Most relevant to this study, fission yeast actin is significantly less efficient than mammalian skeletal muscle actin at activating conventional myosin-II (47). The same behavior has been observed with budding yeast actin (48). Attempts to humanize yeast actin by replacing residues in subdomains 1 and 2 with muscle-specific residues failed to restore robust activation of muscle myosin ATPase activity, suggesting that other parts of the actin molecule must contribute to the actin-myosin interaction (49). This species specificity of myosins and actins has been previously observed for budding yeast myosin-II and myosin-V (50), molecular motors associated with the cytokinetic rings and actin cables, which are composed of formin-nucleated linear actin. *In vitro* studies using purified actin and myosins demonstrated that these motors exhibit greater actin-activated ATPase activity with yeast actin rather than with muscle actin (50). We now provide the first evidence of species specificity for yeast myosin-I, which interacts with a distinct component of the yeast actin cytoskeleton, actin patches composed of branched actin filaments assembled by the Arp2/3 complex. Thus, all myosin classes found in yeast appear to have evolved to work optimally with yeast-specific actin. The inability of HsMyo1e to function in yeast joins the list of other human actin binding proteins that fail to replace budding or fission yeast proteins either due to incompatibility with yeast actin as is the case with cofilin (51) or with yeast orthologs of normal binding partners as reported for profilin (52) and WASP (53).

### Myosin-I protein domain contributions to myosin localization and function

Our work indicates that both head and tail of myosin-I are required for localization at endocytic sites and that each domain makes a distinct contribution to myosin-I function. The fact that HsMyo1e motor domain containing chimeras fail to localize to yeast actin patches suggests that SpMyo1 motor domain is needed not only for the proposed function of generating force to promote endocytic internalization but also for the initial localization of myosin-I at endocytic sites. Conversely, we have found that the HsMyo1e tail is partially functional in yeast when paired with the SpMyo1 motor domain. While these chimeras exhibited enhanced cortical localization compared to SpMyo1, they were recruited to actin patches and partially rescued patch internalization defects in *myo1Δ* cells, indicating that HsMyo1e tail was able to perform at least some of the functions normally carried out by the tail of SpMyo1. The exaggerated membrane localization of the HsMyo1e tail-containing chimeras could be the result of a stronger interaction of HsMyo1e tail with membrane lipids compared to the tail of SpMyo1. It may also result from the lack of myosin regulation, through intra-or intermolecular interactions, which coordinates the timely arrival and departure of SpMyo1 at the membrane during endocytosis. One study using budding yeast has suggested that calmodulin binding regulates autoinhibitory interactions between the CA domain and the neck-TH1 region, with calmodulin dissociation promoting myosin-I localization to actin patches (54). However, we found that in fission yeast, SpMyo1 and all human-yeast myosin-I chimeras recruited Cam1 to patches as long as they contained the SpMyo1 IQ1 motif or HsMyo1e IQ motif, suggesting that the mechanism proposed in budding yeast may not operate in *S. pombe.* Overall, the tail domain of HsMyo1e, when combined with the SpMyo1 motor domain, provides sufficient localization cues to target myosin chimeras to actin patches and support patch internalization.

To determine the roles of individual tail domains, we examined the effects of deletions of the membrane binding TH1 domain and the distal tail, which consists of TH2-SH3 domains in HsMyo1e and TH2-SH3-CA domains in SpMyo1 and is involved in protein-protein interactions. The TH1 domain is thought to act as a lipid anchor positioning myosin so that it can efficiently promote membrane deformation (55). Supporting the role of the TH1 domain as a lipid anchor, a deletional construct of Chimera 3 lacking the HsMyo1e TH1 domain does not localize to actin patches and fails to rescue endocytosis defects in *myo1Δ* cells. Removing the TH1 domain from the full length SpMyo1 also resulted in an inability to rescue endocytosis defects of *myo1Δ* cells, even though SpMyo1ΔTH1, unlike Chimera3ΔTH1, was able to localize to endocytic patches. However, in contrast to SpMyo1 that remains at the base of endocytic invagination during patch internalization, we observed in a portion of actin patches that SpMyo1ΔTH1 internalized with F-actin, indicating a weaker attachment to the cortex. These results are similar to the observations made in *S. cerevisiae* (56), where deletion of the TH1 domain decreased cortical retention of myosin-I and HsMyo1e TH1 domain was able to replace the TH1 domain of yeast myosin-I (57). Thus, in both budding and fission yeast, membrane anchoring by the TH1 domain is critical for myosin-I function in promoting endocytic internalization.

Interestingly, deletion of the TH2-SH3 domains from the HsMyo1e tail of Chimera 3 also results in the loss of localization to patches and the inability to rescue endocytosis in *myo1Δ* cells, suggesting that both the TH1 and TH2-SH3 domains of HsMyo1e are required for endocytic function. This finding agrees with our previous observations, in which deletion of the TH2 domain prevented HsMyo1e localization to sites of HsMyo1e activity in mammalian cells, such as cell-cell junctions or invadopodia (58,59). While the TH2 and SH3 domains are typically involved in protein-protein interactions, the TH2 domain of HsMyo1e, which contains a large number of basic residues, may also contribute to membrane localization (60). The greater contribution of HsMyo1e distal tail to myosin-I localization and function in both yeast and mammalian cells is distinct from the localization requirements of SpMyo1. In agreement with earlier studies (7), deletion of the TH2-SH3-CA domains of SpMyo1 decreases but does not abolish SpMyo1 localization and function at the endocytic sites.

### The utility of yeast system to study myosin-I function

The inability of HsMyo1e motor domain to function in yeast reflects mixed success of attempts to introduce human genes into yeast. Only 47% out of 414 human genes introduced into *S. cerevisiae* were functional due to incompatibility with other subunits within a macromolecular complex or other normal interacting partners (61). This places certain limits on the utility of introducing human genes into yeast both for study of myosin-I function and the effects of disease-associated mutations. Caution is advised when attempting to switch myosin motor domains with different mechanochemical properties as they may not be compatible with yeast actin or may not function properly due to the lack of normal binding partners or actin binding proteins, such as tropomyosins. On the other hand, performing human-yeast domain switching studies using a model system with a straightforward readout, such as yeast endocytosis, provides a solid foundation for future work aiming to characterize the effects of disease-associated mutations in specific myosin domains. In our previous work, the use of the yeast system for functional testing of the effects of kidney disease-associated HsMyo1e mutations was limited to conserved residues in the motor domain of SpMyo1 (21). By demonstrating that chimeras consisting of SpMyo1 motor and HsMyo1e tail functionally complement *myo1Δ* while retaining localization requirements similar to those observed for HsMyo1e in mammalian cells, this study expands the utility of *S. pombe* as a simple model to effectively study disease-associated mutations residing in the HsMyo1e tail.

## EXPERIMENTAL PROCEDURES

### Yeast strain construction

Table S1 lists *S. pombe* strains used in this study. Table S2 lists primers used for strain and plasmid construction. Strains were constructed by homologous recombination-based genomic integrations following lithium acetate transformations (62) and by genetic crosses. To replace *myo1^+^* CDS with *MYO1E* CDS, pBS-SpMyo1 construct, which contains a 5-kb EcoR1 fragment of *S. pombe* genomic DNA encompassing *myo1^+^* gene in pBluescript (7) was linearized by PCR amplification with subM1utr5r and subM1utr3d primers. In parallel, *MYO1E* CDS flanked by 70 nt of *myo1^+^* 5’ UTR and 70 nt of 3’ UTR was amplified from pEGFP-C1-myo1e-EcoRI-, in which EcoRI in *MYO1E* CDS was mutated without changing amino acid sequence. The amplified products were ligated using ClonTech In-Fusion HD Cloning kit (Takara Bio USA, Mountain View, CA) and the resultant pBS-HsMyo1e construct contained *MYO1E* CDS flanked by *myo1^+^* 5’ and 3’ UTRs. The HsMyo1e E337S TEDS site mutation was generated in pBS-HsMyo1e construct using the QuickChange kit (Stratagene; La Jolla, CA) according to the manufacturer’s instructions. To replace *S. pombe myo1^+^* CDS with coding sequences for Myo1e or Myo1e(E337S) in the endogenous *myo1* locus, the pBS constructs were digested with EcoR1, and 5-kb fragments containing HsMyo1e CDS or HsMyo1e(E337S) CDS flanked by *myo1^+^* 5’ and 3’ UTRs were introduced via lithium acetate transformation into the *myo1Δ ::ura4^+^* strain (TP192), in which the *myo1^+^* CDS was replaced with the *ura4^+^* nutritional marker (13), followed by counter-selection against *ura4^+^* on EMM agar plates containing 2 mg/ml 5-FOA (5-fluoroorotic acid) and 0.05 mg/ml uracil. Positive clones were identified by PCR and verified by sequencing. These strains express untagged Myo1e and Myo1e(E337S) from the native *myo1* locus under control of the endogenous *Pmyo1* promoter. To tag HsMyo1e and HsMyo1e(E337S) expressed from *myo1* locus with mGFP at the C-terminus, the stop codon in the HsMyo1e coding sequence was replaced with mGFP-tagging cassette amplified from pFA6a-mGFP-kanMX6 (pJQW85-4) using primers m1e-GFP-int-F and m1-Kxr (63,64). The integrations were confirmed by PCR and sequencing.

Strains with different combinations of gene deletions and fluorescent protein-tagged alleles were generated by genetic crosses. Cells were mated on ME (Malt Extract) agar plates, followed by tetrad dissections on YES agar plates and screening for wanted gene combinations by replica plating onto appropriate selective plates, microscopy, and PCR diagnostics. To facilitate mating, in some cases *myo1Δ* cells were transformed with pUR-myo1 plasmid (7), which is subsequently lost upon sporulation.

### Plasmid construction

Plasmids for expression of mGFP-tagged myosin constructs in *S. pombe* are listed in Table S3. For the purposes of making chimeras, *S. pombe* Myo1 domains were defined as follows: motor domain, aa 1-720; IQ1, aa 721-741; IQ2, aa 742-763; TH1, aa 764-966; TH2, aa 967-1112; SH3, aa 1113-1163; CA, 1164-1217. *H. sapiens* Myo1e domain boundaries were similarly defined: motor domain, aa 1-692; IQ, aa 693-717; TH1, aa 718-923; TH2, aa 924-1055; SH3, aa 1056-1108.

Chimeras 1-5 were first generated in pBluescript by In-Fusion cloning (Takara Bio, USA, Mountain View, CA) of inserts encoding different fragments of human Myo1e PCR-amplified from pEGFP-C1-myo1e-EcoR1-into the PCR-amplified segments of pBS-SpMyo1 vector using primers designed to replace selected SpMyo1 domain sequences with corresponding HsMyo1e sequences. For Chimera 1, a fragment encoding the HsMyo1e motor and IQ domains was amplified with primers m1Myo1Ed and SpHs1hA and ligated into pBS-SpMyo1 amplified with primers subM1utr5r and SpHs1p1, thereby replacing *S. pombe* Myo1 motor, IQ1 and IQ2 domains. For Chimera 2, a fragment encoding HsMyo1e motor domain was amplified with primers m1Myo1Ed and SpHs2hB and ligated into pBS-SpMyo1 amplified with primers subM1utr5r and SpHs2p2 thereby replacing *S. pombe* Myo1 motor domain. For Chimera 5, a fragment encoding HsMyo1e motor and IQ domains was amplified with primers m1Myo1Ed and SpHs5hE and ligated into pBS-SpMyo1 amplified with primers subM1utr5r and SpHs5p5 thereby replacing *S. pombe* Myo1 motor and IQ1 domains. For Chimera 3, a fragment encoding HsMyo1e tail consisting of the TH1, TH2, and SH3 domains was amplified with primers SpHs3hC and m1Myo1Er and ligated into pBS-SpMyo1 amplified with primers SpHs3p3 and subM1utr3d thereby replacing *S. pombe* Myo1 tail consisting of TH1, TH2, SH3, and CA domains. For Chimera 4, a fragment encoding HsMyo1e IQ and tail domains was amplified with primers SpHs4hD and m1Myo1Er and ligated into pBS-SpMyo1 amplified with primers SpHs4p4 and subM1utr3d thereby replacing *S. pombe* Myo1 IQ1, IQ2 and tail domains.

To move into mGFP-tagging vector for yeast expression, SpMyo1, HsMyo1e, HsMyo1e(E337S), and Chimeras 1-5 sequences were amplified using the appropriate primer pairs (selected from human primers Myo1e F, Myo1e R and yeast primers Myo1 F, SpTail R) containing NotI and NheI cloning sites, subcloned into pCR-BluntII-TOPO using the Zero Blunt TOPO PCR Cloning Kit (Thermo Fisher Scientific, Waltham, MA) and verified by sequencing. The inserts were then cut from the TOPO constructs using NotI and NheI and ligated into the pSGP-573 vector (65) that we previously modified by replacing the GFP sequence with monomeric mGFP and adding GASGTGS linker at the mGFP C-terminus. All constructs in this modified pSGP-573 vector express N-terminally mGFP-tagged proteins under control of the strong thiamine-repressible *3xPnmt1* promoter (66). To generate Chimera 6, the IQ2 domain was removed from pSGP-573-Chimera 3 by linearizing the plasmid using primers Chimera6 and m1D1IQ2L followed by ligation using In-Fusion cloning. The HsMyo1e tail construct was derived from pSGP-573-HsMyo1e using primers pSPG-m1etail-F and pSPG-m1etail-R followed by ligation using In-Fusion cloning. The SpMyo1ΔTH1 and SpMyo1 ΔTH2-SH3-CA constructs were derived from pSGP-573-SpMyo1 by amplifying with primers SpMyo1deltaTH1 and IQ2rev to remove TH1 domain coding sequence and primers SpMyo1delta23CA and SpMyo1TH1rev to remove TH2-SH3-CA coding sequence followed by ligation using In-Fusion cloning. The Chimera 3ΔTH1 and Chimera 3ΔTH2-SH3 constructs were generated from pSGP-573-Chimera 3 by amplifying with primers CHIM3deltaTH1 and IQ2rev to remove TH1 domain coding sequence and primers Sp1m-m1e-TH1-F and hmyo1edelTH2 to remove TH2-SH3 coding sequence followed by ligation using In-Fusion cloning. Empty modified pSGP-573 vector expressing mGFP alone was used as a negative control.

### Yeast electroporation

For electroporation, cells were grown overnight in EMMS media to an OD_595_ of 0.4-1.5, pelleted at 3,000 rpm in Beckman (Brea, CA) Allegra X-15R centrifuge at 4°C, washed twice with cold sterile water, once with cold sterile 1M sorbitol, and re-suspended in sorbitol to a concentration of ~2 x 10^9^ cells/mL. Each plasmid was added to 40μL of cells and incubated on ice for 7 minutes. Cells were then electroporated using the Bio-Rad (Hercules, CA) Micropulser Electroporator according to the instruction manual, plated onto EMM agar plates lacking uracil and containing 5 μg/ml thiamine, and incubated at 25°C.

### Salt sensitivity assays

Stably integrated strains were streaked onto YES agar plates containing 1M KCl and incubated for several days at 25°C. Cells transformed with constructs in pSGP-573 vector were streaked onto EMM agar plates lacking uracil and containing 1M KCl with or without 5 μg/ml thiamine.

### Fluorescence microscopy

Stably integrated strains were grown to exponential phase in EMM5S medium at 25°C over two days. Fission yeast cells transformed with constructs in pSGP-573 vector were grown to exponential phase over two days at 25°C in EMM medium lacking uracil (EMM-ura) and containing 5 μg/ml thiamine. Cells were then washed 3 times in EMM-ura, seeded at OD_595_=0.08 into in EMM-ura without thiamine, and incubated for 12-26 hours to induce protein expression. Generally, constructs containing the HsMyo1e motor domain needed longer induction times.

For imaging, cells were pelleted briefly using a tabletop centrifuge and mounted on pads of 25% gelatin in EMM5S under coverslips sealed with Valap. Following 5 minutes of equilibration, cells were imaged at ambient room temperature using a PerkinElmer (Waltham, MA) UltraView VoX Spinning Disc Confocal system mounted on a Nikon Eclipse Ti-E microscope equipped with Hamamatsu C9100-50 EMCCD camera and a 100×/1.4 N.A. PlanApo objective, controlled by Volocity software. For Z-series through the entire cells, images were captured at 0.4-μm intervals. Time series of 2-color images were collected in the middle plane of the cells at 2-second intervals for 1 minute.

### Image analysis

All image analysis was performed using Fiji (National Institutes of Health, Bethesda, MD)(67). Whole-cell fluorescence intensities of mGFP-tagged proteins in individual cells, which represent protein expression levels, were averaged from frames 4-8 of a time series at the mid-cell section and subtracted for extracellular background. Only cells expressing exogenous myosin within 0.5–2-fold of the level of mGFP-SpMyo1 expressed from the native *myo1* locus under control of the endogenous *Pmyo1* promoter (13) were analyzed for tracking patch intensities and movement, percent internalization, and protein colocalization. For actin patch tracking, at least 10 patches from 3-5 cells were analyzed for each strain. Using a 10-pixel wide circular ROI, the patch fluorescence intensity and position were tracked for both myosin and Fim1-mCherry over the duration of each patch lifetime. Individual time courses of patch intensity and distance from the origin were aligned to the peak intensity of Fim1-mCherry at time zero and averaged at each time point. Percent internalization was measured by following patches over a 20-second time period in at least 8 cells. Internalization was considered successful if the patch moved at least 2 pixels (130 nm) from the cell membrane. Colocalization between mGFP-tagged myosin constructs and mCherry-tagged fimbrin Fim1 or calmodulin Cam1 in patches was measured over a 20-second time period by using the Plot Z function to detect the presence of mGFP and mCherry signals within a 10-pixel wide ROI. At least 20 patches in 5-10 cells were analyzed for each strain. All strains were imaged at least twice to assess reproducibility.

### Western blots

Cell cultures grown for live cell imaging or grown under the same conditions as for live cell imaging were harvested by centrifugation for 5 minutes at 3,000 rpm in Beckman (Brea, CA) Allegra X-15R centrifuge at 4°C and processed for immunoblotting as previously described (68). Pelleted cells were re-suspended in ice-cold lysis buffer U (50 mM HEPES pH7.5, 100 mM KCl, 3 mM MgCl2, 1 mM EGTA, 1 mM EDTA, 0.1% Triton X-100, 1 mM DTT, 1 mM PMSF and Complete (Roche, Branchburg, NJ) protease inhibitor cocktail), mixed with glass beads and lysed mechanically using the FastPrep-24 (MP-Bio, Santa Ana, CA), followed by addition of SDS-PAGE sample buffer. Lysates were then heated for 2 minutes at 100°C, followed by a 5 minute centrifugation at 7,000 rpm. Total protein samples were adjusted based on an OD_595_ reading prior to centrifugation, separated on 10-20% gradient SDS-PAGE gels, and transferred to Immobilon-P (EMD Millipore, Billerica, MA). Membranes were exposed to primary anti-GFP antibody (67 MA5-15256, Pierce, Rockford, IL) at 1:3000 and incubated overnight at 4°C. The next day, the blot was washed with TBS-T 3 times for 5 minutes and incubated with secondary horseradish-peroxidase-conjugated antibody for 1 hour at room temperature. Blots were developed using Clarity™ Western ECL substrate (Bio-Rad, Hercules, CA) and imaged using a Bio-Rad (Hercules, CA) ChemiDoc MP imager.

### Statistical analysis

Graphpad Prism Software was used for graphing, linear regression and all other statistical analysis. For multiple comparisons, data were analyzed using a using a one-way ANOVA with Tukey’s Post-Hoc test, with statistical significance set at a p-value < 0.05.

## ACKNOWLEDGMENTS

This work was supported by an AHA (18PRE34070066) fellowship to S.R.B., AHA SDG 11SDG5470024 to V.S. and the National Institute of Diabetes and Digestive and Kidney Diseases of the NIH under Award R01DK083345 to M.K. We thank Robert Carroll and Cameron MacQuarrie for helpful discussions. The content is solely the responsibility of the authors and does not necessarily represent the official views of the National Institutes of Health.

## CONFLICT OF INTEREST

The authors declare that they have no conflicts of interest with the contents of this article.

## ABBREVIATIONS AND NOMENCLATURE

The abbreviations used are: Myo1e, myosin 1e; HsMyo1e, human myosin 1e; SpMyo1, *S. pombe* myosin-I; Fim1, fimbrin 1; Cam1, calmodulin; Cam2, calmodulin-related light chain 2; TH1, tail homology 1; TH2, tail homology 2; SH3, Src homology 3; CA, central-acidic

## SUPPORTING INFORMATION

**Figure S1:**
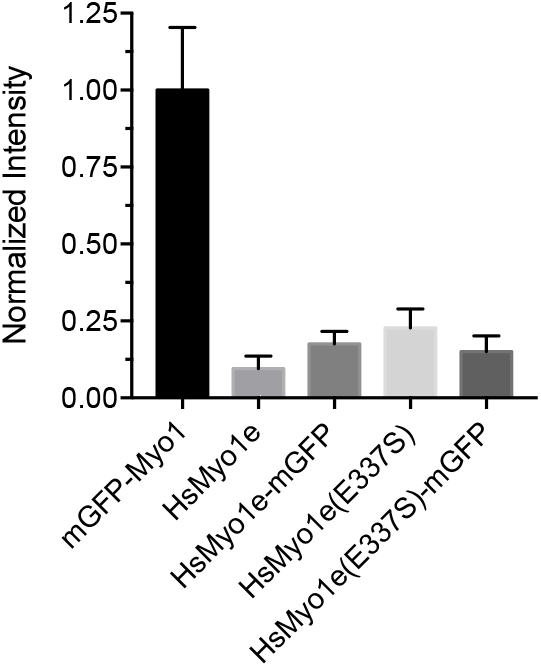
HsMyo1e fails to express from endogenous *S. pombe myo1* locus. Bar graph (mean±SD) of whole cell fluorescence intensities of *S. pombe* cells expressing from the *myo1* locus under control of endogenous *Pmyo1* promoter mGFP-SpMyo1 (mGFP-Myo1), untagged HsMyo1e, mGFP-tagged HsMyo1e, untagged HsMyo1e(E337S), or mGFP-tagged HsMyo1e(E337S). Fluorescence intensities were measured in a single confocal section through the middle of the cells, subtracted for extracellular background, and normalized to the intensity of mGFP-Myo1. Untagged strains were used as controls for autofluorescence. N = 18-20 cells.

**Figure S2:**
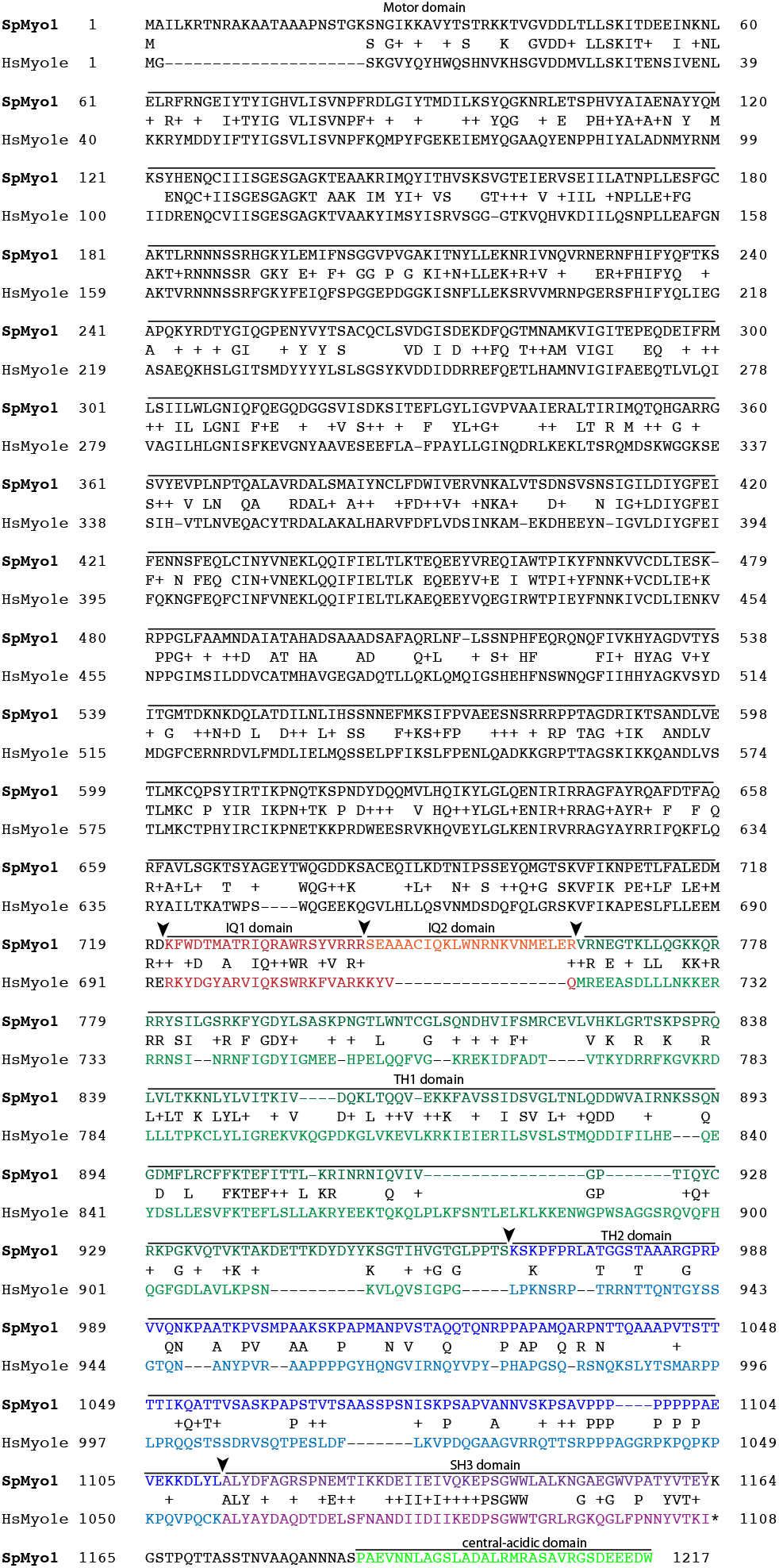
Alignment of *S. pombe* Myo1 (SpMyo1) and *H. sapiens* Myo1e (HsMyo1e) protein sequences. Protein domains of SpMyo1 and HsMyo1e are color coded and labeled: motor domain (black), IQ1 (red), IQ2 (orange), TH1 (green), TH2 (blue), SH3 (purple), CA (bright green). Identical and similar amino acid residues are indicated on the middle consensus line. Black arrowheads mark domain boundaries used to create chimeric and deletion constructs. TH, tail homology domain; SH3, Src Homology 3 domain; CA, central-acidic domain.

**Figure S3:**
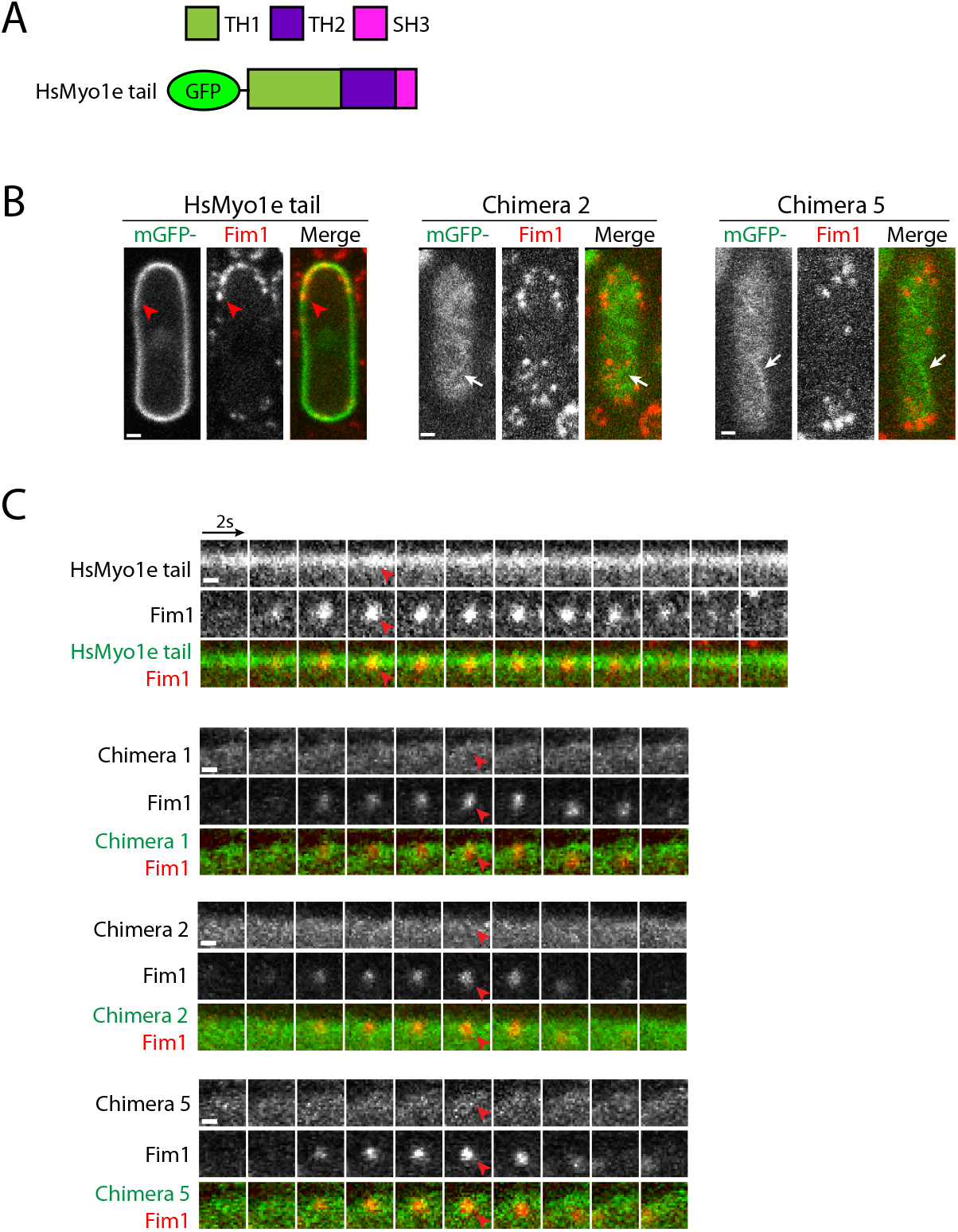
HsMyo1e tail and HsMyo1e motor – SpMyo1 tail chimeras do not localize to actin patches. (A-C) Analysis of colocalization of mGFP-tagged (green) HsMyo1e tail or HsMyo1e motor - SpMyo1 tail chimeras 1, 2, and 5 with Fim1-mCherry (red) in actin patches in *myo1Δ* cells. mGFP-tagged constructs were expressed from plasmids under control of *3xPnmt1* promoter for 12-18 hours in the absence of thiamine in *myo1Δ* cells expressing Fim1-mCherry. (A) Schematic diagram of mGFP-tagged HsMyo1e tail construct. (B) Representative images in single confocal sections through the middle (HsMyo1e tail) or the top surface (Chimeras 2 and 5) of the cells. Scale bars, 1 μm. (C) Montages of individual patches at 2-second intervals. Scale bar, 0.5 μm. The white arrows indicate Chimeras 2 and 5 in the filamentous thread-like structures on the surface of the cell. Red arrowheads indicate the absence of mGFP-tagged proteins in Fim1-mCherry-labeled actin patches.

**Figure S4:**
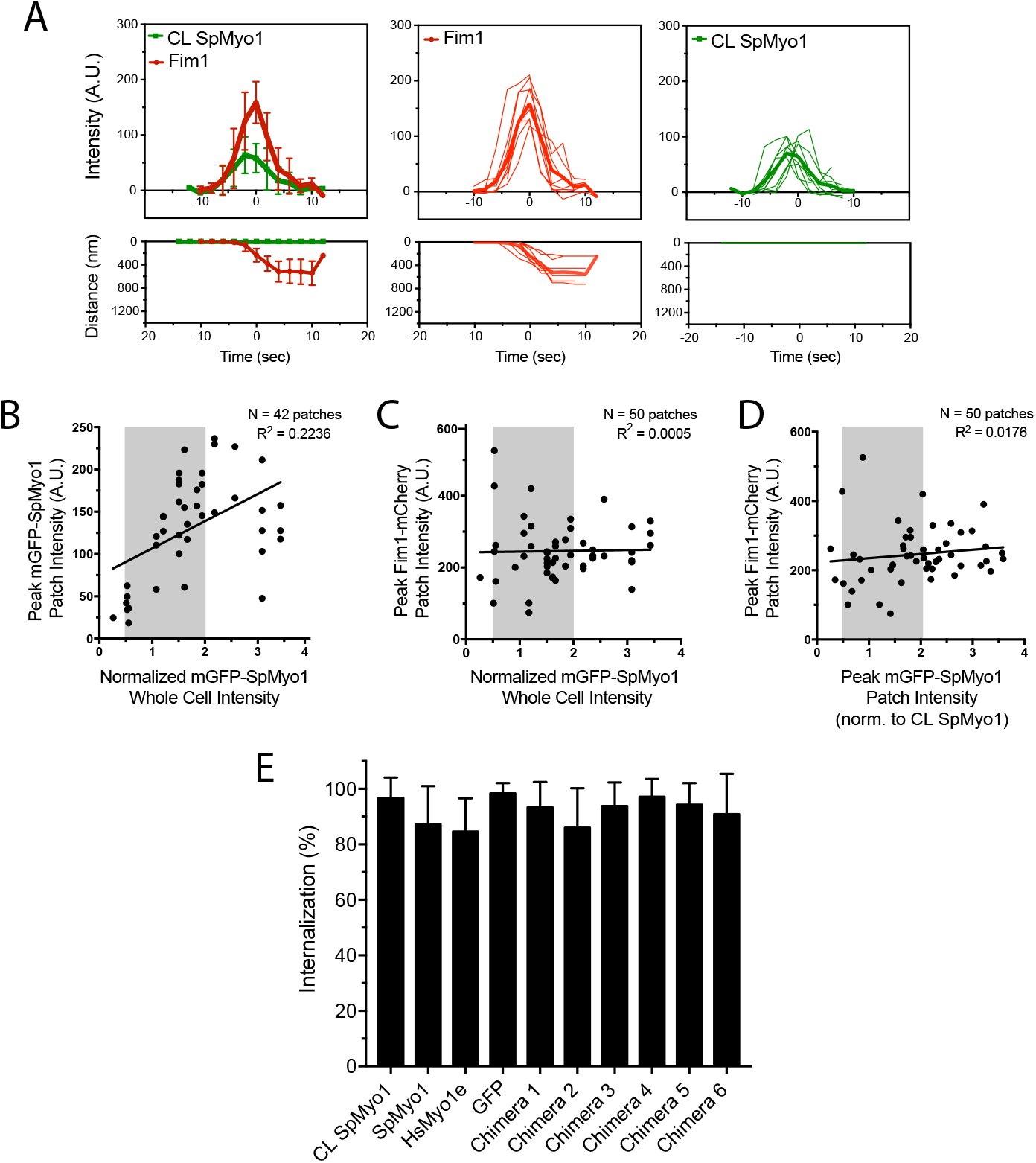
Tracking dynamics of mGFP-tagged SpMyo1 and human-yeast myosin-I chimeras, and Fim1-mCherry in endocytic actin patches. (A) Raw (thin lines) and average (thick lines; error bars in the graphs on the left indicate SD) time courses of (upper panels) fluorescence intensity and (lower panels) distance traveled for mGFP-Myo1 (green) and Fim1-mCherry (red) in actin patches in control wild-type cells expressing mGFP-Myo1 (CL SpMyo1) from the endogenous *myo1* locus. Patch dynamics were tracked in time series of images acquired at 2-second intervals in a single confocal section through the middle of the cells. The time courses of cortical background-subtracted intensities and distances from the origin for individual patches were aligned to the peak of Fim1-mCherry patch intensity (time zero) and averaged at each time point. N= 9 patches in 3 cells. (B-D) Correlation plots of (B) peak intensities of mGFP-SpMyo1 in patches versus whole cell mGFP-SpMyo1 intensities, (C) peak intensities of Fim1-mCherry in patches versus whole cell mGFP-SpMyo1 intensities, and (D) Fim1-mCherry peak patch intensities versus mGFP-SpMyo1 peak patch intensities. mGFP-SpMyo1 was expressed from the plasmid under control of *3xPnmt1* promoter for 12 hours in the absence of thiamine in *myo1Δ* cells expressing Fim1-mCherry. Whole cell intensities and peak patch intensities were measured in time series of images acquired at 2-second intervals in a single confocal section through the middle of the cells. Whole cell intensities representing mGFP-SpMyo1 expression levels were normalized to the intensities of control wild-type cells expressing mGFP-Myo1 from the endogenous *myo1* locus. Lines represent the best linear fits with corresponding R^2^ values from linear regression analysis. N=42-50 patches from 20 cells. Shaded areas indicate the 0.5-2-fold range of expression levels that was accepted for tracking dynamics of myosin and Fim1 in actin patches. (E) Bar graph of percent internalization (±SD) of Fim1-mCherry patches in wild-type cells expressing mGFP-tagged SpMyo1, SpMyo1e, or human-yeast myosin-I chimeras from the plasmid under control of *3xPnmt1* promoter for 12-16 hours in the absence of thiamine. N= 28-45 patches in at least 5 cells. No significant differences from wild-type cells (CL SpMyo1) expressing mGFP-SpMyo1 from the endogenous *myo1* locus were determined by a one-way ANOVA (p=0.3839).

**Figure S5:**
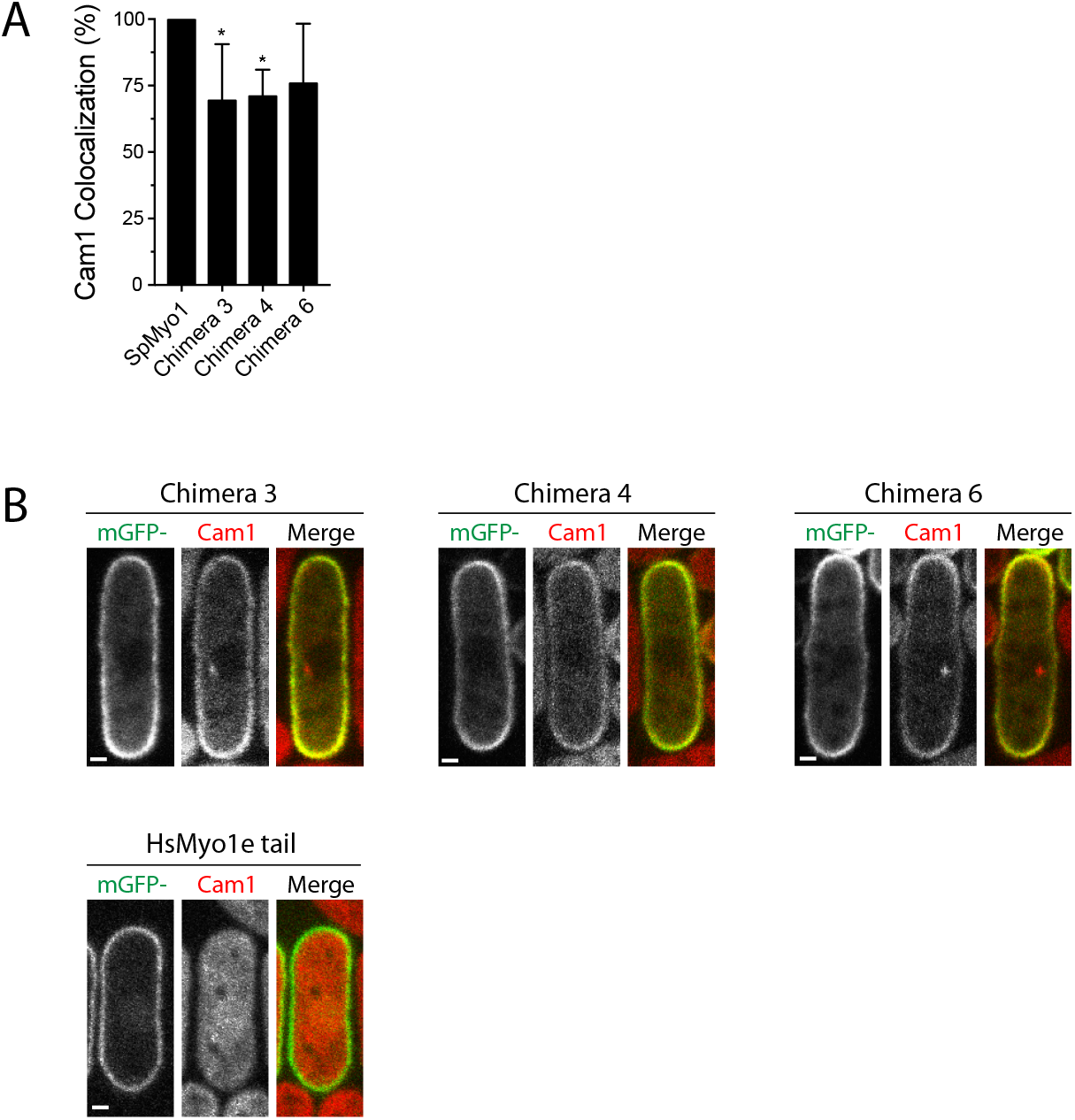
SpMyo1 motor – HsMyo1e tail chimeras localize to the cell cortex and recruit Cam1. (A) Bar graph of the percent colocalization (±SD) of Cam1 and SpMyo1 or SpMyo1 motor - HsMyo1e tail chimeras in cortical actin patches. N=24 patches in 6 cells. Asterisks indicate a significant difference from SpMyo1 as determined by a one-way ANOVA, * p<0.05. (B) Images in a single confocal section through the middle of the *myo1Δ* cells expressing mCherry-Cam1 (red) and overexpressing mGFP-tagged (green) SpMyo1 motor - HsMyo1e tail chimeras 3,4, and 6 or HsMyo1e tail. mGFP tagged constructs were expressed from plasmids under control of *3xPnmt1* promoter for 13-18 hours in the absence of thiamine. Scale bars, 1μm.

**Figure S6:**
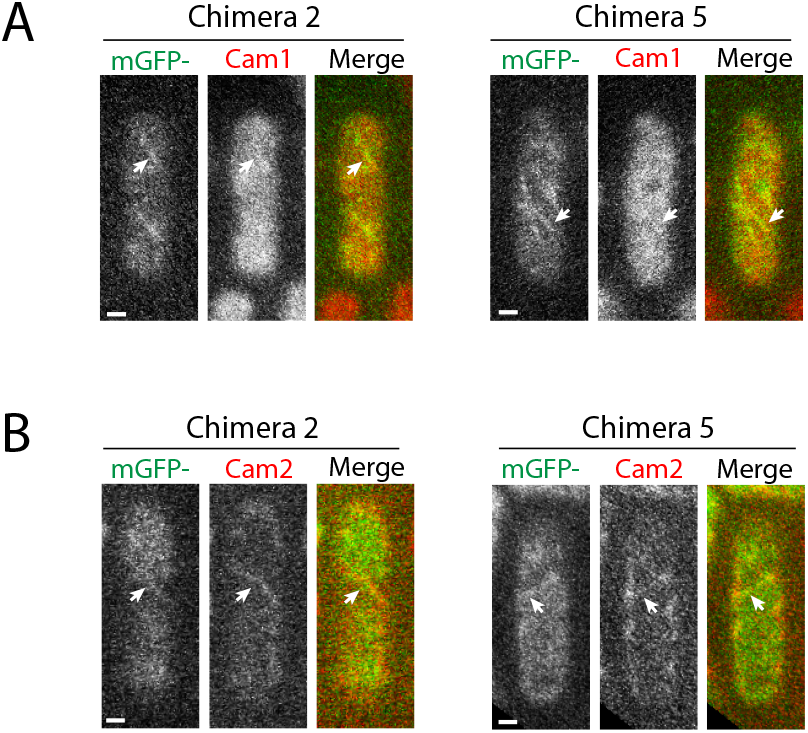
HsMyo1e motor - SpMyo1 tail Chimeras 2 and 5 localize to cortical filamentous structures and recruit Cam1 and Cam 2. (A, B) Images in a single confocal section through the top surface of *myo1Δ* cells expressing mGFP-tagged (green) HsMyo1e motor - SpMyo1 tail chimeras 2 and 5 from plasmids under control of *3xPnmt1* promoter and also expressing (A) mCherry-Cam1 or (B) Cam2-mCherry (red). White arrows indicate colocalization of Cam1 or Cam2 with HsMyo1e motor - SpMyo1 tail chimeras in filamentous thread-like cortical structures. Scale bars, 1 μm.

**Figure S7:**
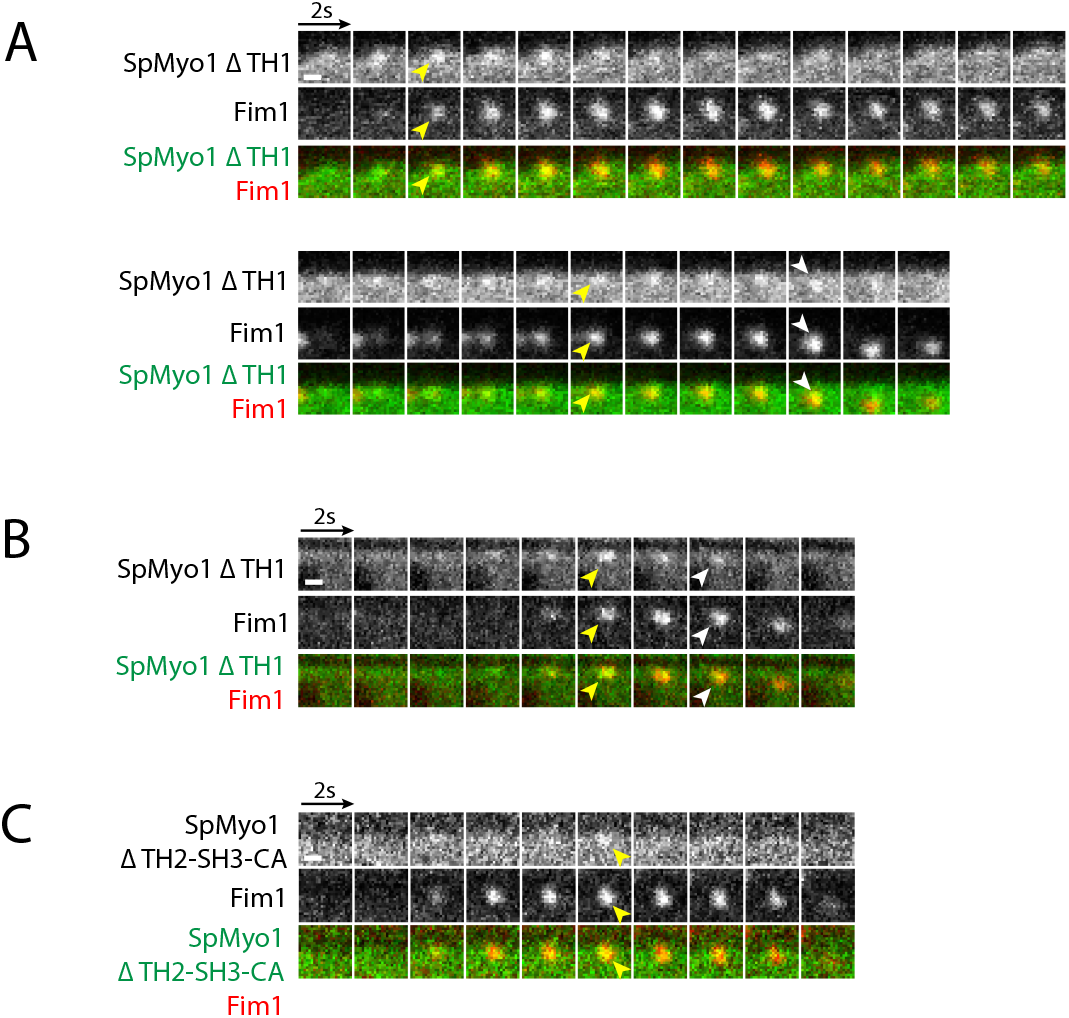
SpMyo1 does not require TH1 or TH2-SH3-CA domains for localization to endocytic actin patches. (A, B) Example montages of non-internalizing (top panel) and internalizing (bottom panel) patches in *myo1Δ* cells expressing mGFP-tagged (green) SpMyo1ΔTH1 and Fim1-mCherry (red). (B) An example of an internalizing patch in wild-type cells expressing mGFP-tagged (green) SpMyo1ΔTH1 and Fim1-mCherry (red). (C) An example of an internalizing patch in wild-type cells expressing mGFP-tagged (green) SpMyo1ΔTH2-SH3-CA and Fim1-mCherry (red). mGFP-tagged constructs were expressed from plasmids under control of *3xPnmt1* promoter for 12-15 hours in the absence of thiamine. All montages are presented at 2-second intervals in a single confocal section through the middle of the cell. Yellow arrowheads indicate colocalization of myosin constructs and Fim1-mCherry in actin patches. White arrows depict internalization of mGFP-SpMyo1ΔTH1 with Fim1-mCherry, Scale bar, 0.5 μm.

**Table S1.**
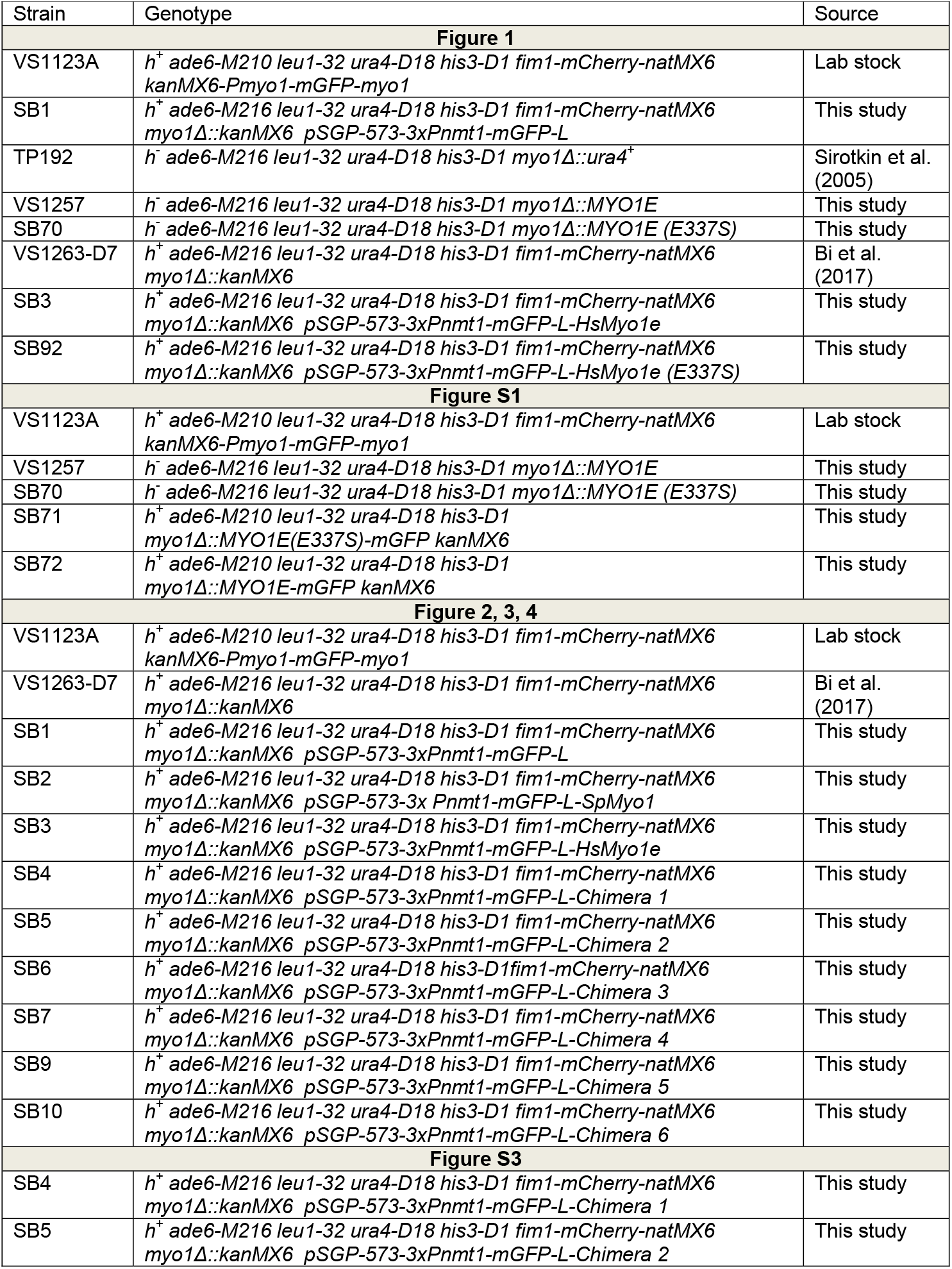
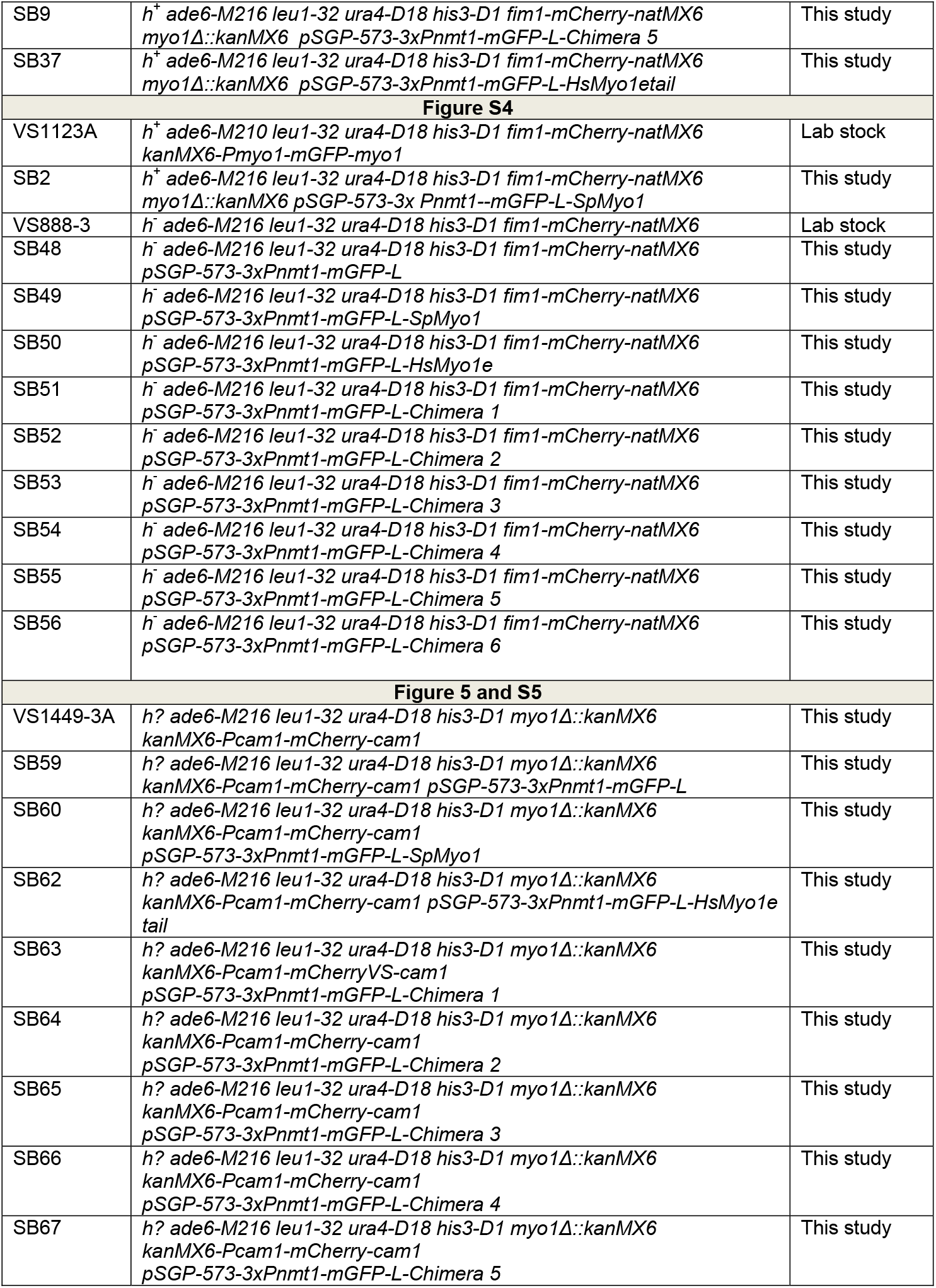
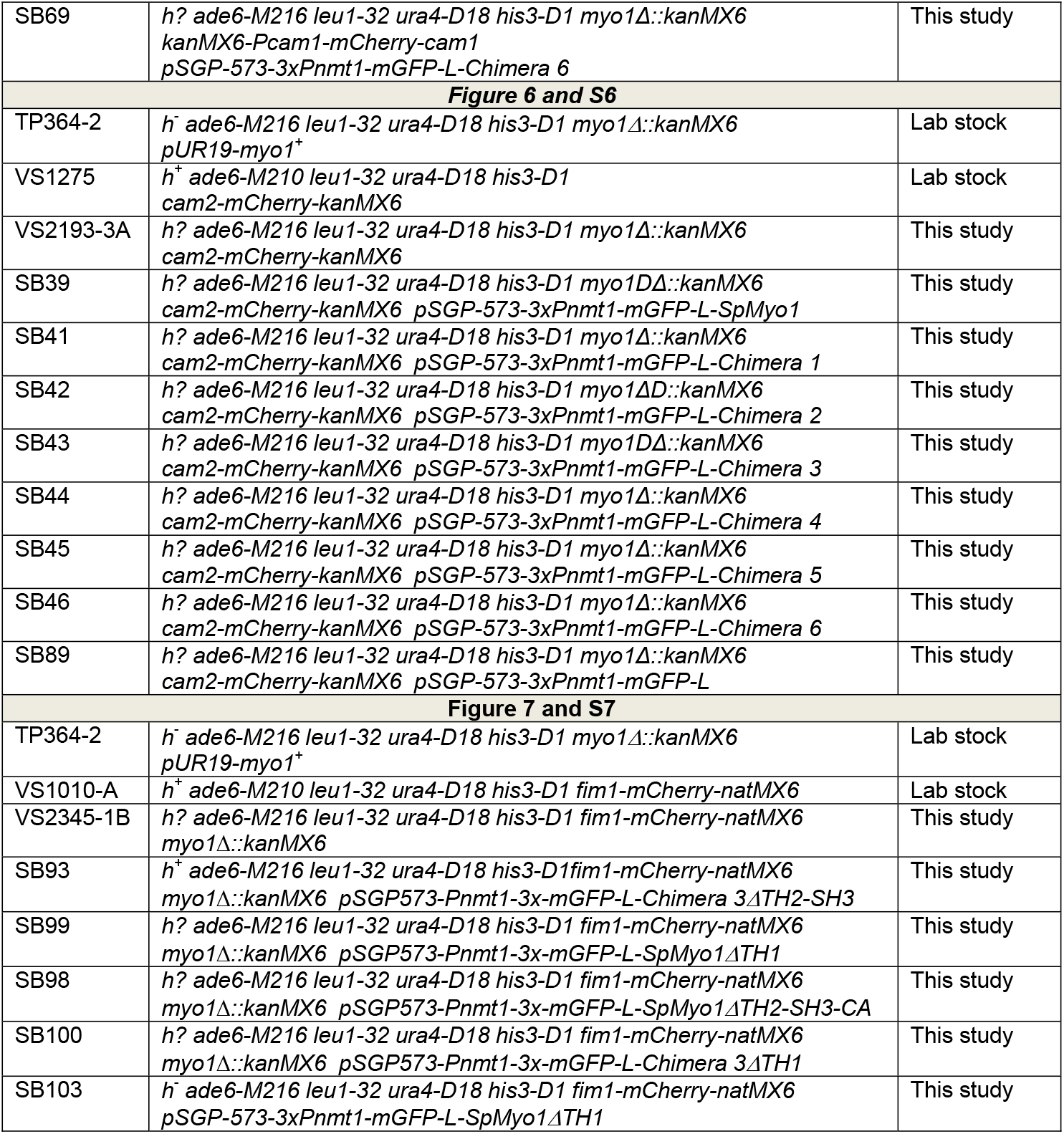
*S. pombe* strains used in this study.

**Table S2.**
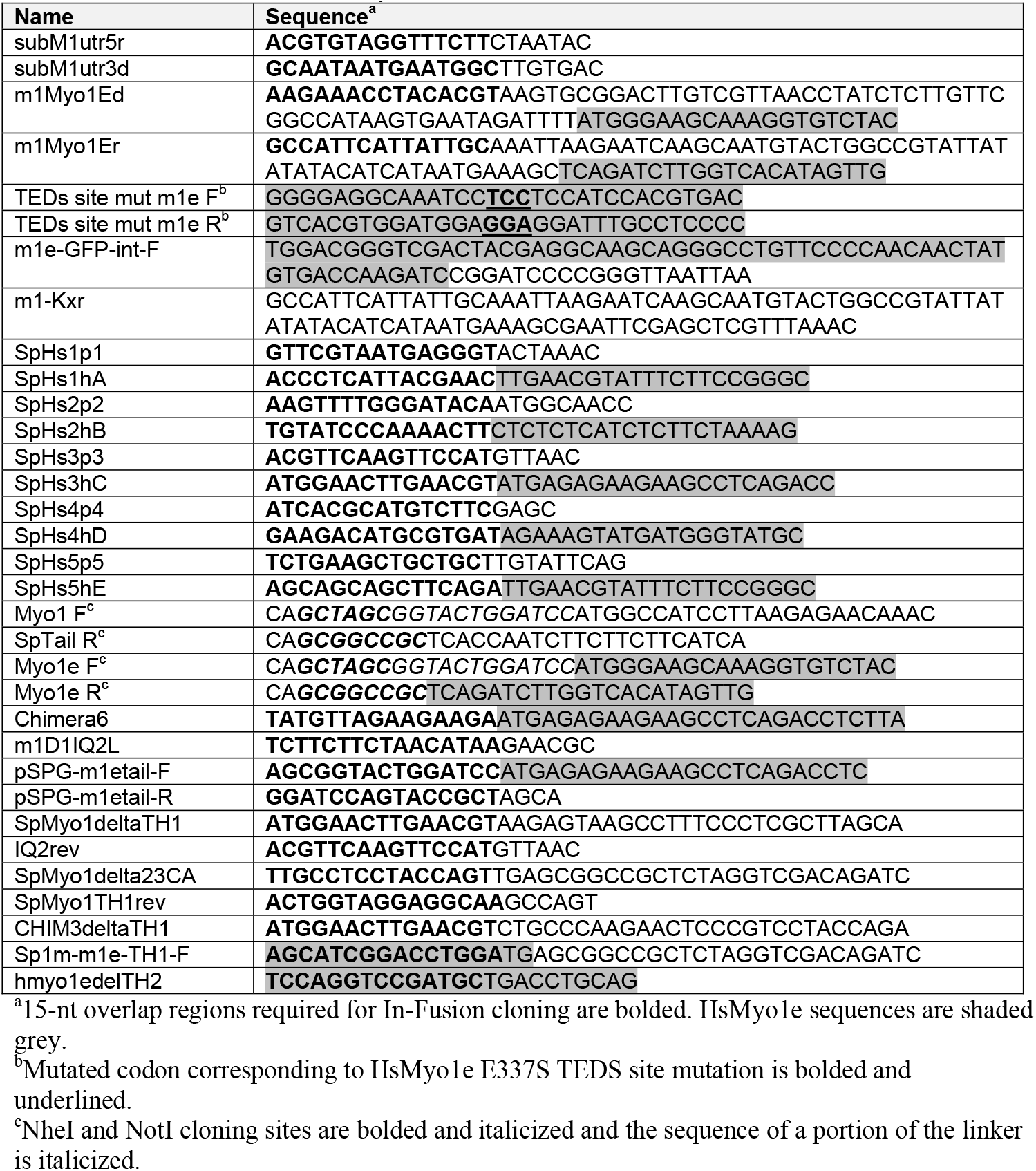
Primers used in this study.

**Table S3.** Plasmids for expression of mGFP-tagged proteins used in this study and templates and primers used for their construction.

| **Construct<sup>a</sup>** | **Description** | **Hs template** |  | **Sp template** |  | **Subcloning primers** |
| --- | --- | --- | --- | --- | --- | --- |
| SpMyo1 | *S. pombe* Myo1 |  |  | pBS-SpMyo1 |  | Myo1 F<br>SpTail R |
| HsMyo1e | Human Myo1e | pBS-HsMyo1e |  |  |  | Myo1e F<br>Myo1e R |
| Hs Myo1e (E337S) | Human Myo1e TEDS site mutant | pBS-HsMyo1e (E337S) |  |  |  | Myo1e F<br>Myo1e R |
| **Construct<sup>a</sup>** | **Description** | **Hs template** | **Hs primers** | **Sp template** | **Sp primers** | **Subcloning primers** |
| Chimera 1 | HsMyo1e motor-IQ<br>SpMyo1 tail | pEGFP-C1-<br>myo1e-EcoR1- | m1Myo1Ed<br>SpHs1hA | pBS-SpMyo1 | subM1utr5r<br>SpHs1p1 | Myo1e F<br>SpTail R |
| Chimera 2 | HsMyo1e motor<br>SpMyo1 IQ1-IQ2-tail | pEGFP-C1-<br>myo1e-EcoR1- | m1Myo1Ed<br>SpHs2hB | pBS-SpMyo1 | subM1utr5r<br>SpHs2p2 | Myo1e F<br>SpTail R |
| Chimera 3 | SpMyo1 motor-IQ1-IQ2<br>HsMyo1e tail | pEGFP-C1-<br>myo1e-EcoR1- | SpHs3hC<br>m1Myo1Er | pBS-SpMyo1 | SpHs3p3<br>subM1utr3d | Myo1 F<br>Myo1e R |
| Chimera 4 | SpMyo1 motor<br>HsMyo1e IQ-tail | pEGFP-C1-<br>myo1e-EcoR1- | SpHs4hD<br>m1Myo1Er | pBS-SpMyo1 | SpHs4p4<br>subM1utr3d | Myo1 F<br>Myo1e R |
| Chimera 5 | HsMyo1e motor-IQ<br>SpMyo1 IQ2-tail | pEGFP-C1-<br>myo1e-EcoR1- | m1Myo1Ed<br>SpHs5hE | pBS-SpMyo1 | subM1utr5r<br>SpHs5p5 | Myo1e F<br>SpTail R |
| **Construct<sup>b</sup>** | **Description** | **Template** | **Primers** |  |  |  |
| Chimera 6 | SpMyo1 motor-IQ1<br>HsMyo1e tail | Chimera 3 | Chimera6<br>m1D1IQ2L |  |  |  |
| HsMyo1e tail | HsMyo1e tail | HsMyo1e | pSPG-m1etail-F<br>pSPG-m1etail-R |  |  |  |
| SpMyo1ΔTH1 | SpMyo1ΔTH1 | SpMyo1 | SpMyo1deltaTH1<br>IQ2rev | | | |
| SpMyo1ΔTH2-SH3-CA | SpMyo1ΔTH2-SH3-CA | SpMyo1 | SpMyo1delta23CA<br>SpMyo1TH1rev | | | |
| Chimera 3ΔTH1 | Chimera 3ΔTH1 | Chimera 3 | CHIM3deltaTH1<br>IQ2rev | | | |
| Chimera 3ΔTH2-SH3 | Chimera 3ΔTH2-SH3 | Chimera 3 | Sp1m-m1e-TH1-F<br>hmyo1edelTH2 | | | |
| GFP | Modified pSGP-573 |  |  |  |  |  |

<sup>a</sup>SpMyo1, HsMyo1e, HsMyo1e (E337E), and Chimeras 1-5 were first made in pBluescript, then amplified by PCR and cloned into pCR-BluntII-TOPO followed by subcloning into modified pSGP-573 vector, in which GFP sequence was replaced with mGFP sequence and N-terminal mGFP is separated from C-terminal portion of the fusion protein by 7-aa GASGTGS linker.
<sup>b</sup>Chimera 6 was derived from Chimera 3 by In-Fusion cloning. HsMyo1e tail construct was similarly derived from HsMyo1e construct. The ΔTH1 and ΔTH2-SH3-CA mutants of SpMyo1 and the ΔTH1 and ΔTH2-SH3 mutants of Chimera3 were derived by In-Fusion cloning from SpMyo1 and Chimera 3 constructs, respectively.

